# Beyond Immune Evasion: CD47-Driven Pro-Tumorigenic Dysfunction in Diffuse Large B Cell Lymphoma and Triple-Negative Breast Cancer

**DOI:** 10.64898/2026.07.25.740652

**Authors:** Travis Cheng In Lum, Jordan Yong Ming Tan, Felix Jun Heng Ng, Sai Mun Leong, Muhammad Sufyan Bin Masroni, Susan Swee Shan Hue

**Author notes:** These authors contributed equally to this work (Travis Cheng In Lum, Jordan Yong Ming Tan). Senior corresponding author (Susan Swee Shan Hue).

## Abstract

CD47 is a ubiquitously expressed transmembrane protein that functions as a negative immune checkpoint, marking host cells as “self” by delivering an inhibitory “don’t-eat-me” signal to phagocytes. Cancer cells co-opt this mechanism, upregulating CD47 to evade immunosurveillance and phagocytosis by innate immune cells, a pattern observed across solid tumours and haematological malignancies. CD47 overexpression correlates with poor prognosis across most cancer types, including therapy-resistant disease.

Despite extensive efforts to develop CD47-targeted therapies, the downstream biological consequences of aberrant CD47 expression within tumour cells remain poorly characterised beyond its established anti-phagocytic role. This study investigated non-immunological, pro-tumorigenic functions of CD47 to define the cellular effects, beyond immune evasion, that CD47-targeted therapy might disrupt. We found that CD47 exerts cancer type-specific effects: in DLBCL, CD47 loss impaired mitochondrial metabolism and sensitised cells to R-CHOP standard-of-care chemoimmunotherapy, whereas in triple-negative breast cancer (TNBC), CD47 knockdown delayed cell cycle progression, enhanced migration, and conferred resistance to specific chemotherapeutic agents.

These findings indicate that CD47 has multifaceted, context-dependent roles in tumour biology that extend beyond immune checkpoint signalling. Clinically, this suggests CD47-targeted therapies may produce cancer type-specific off-target effects on tumour metabolism, proliferation, and drug sensitivity, which are considerations that should inform their rational combination with existing targeted therapies.

## 1. Introduction

### 1.1 Diffuse Large B-Cell Lymphoma

Diffuse large B-cell lymphoma (DLBCL) is one of the most common hematological cancers worldwide, accounting for approximately 30–60% of all non-Hodgkin lymphomas (NHL)^1^. As an aggressive NHL subtype, DLBCL displays substantial disease heterogeneity, reflected in its distinct molecular (morphology, genetics), pathological, and clinical features. DLBCL arises from the malignant transformation of mature B cells at various stages of differentiation during the germinal center (GC) reaction, a dynamic microanatomical response that develops in secondary lymphoid organs upon antigen exposure^2^. As the principal site of clonal expansion and antibody affinity maturation, the GC comprises two functionally distinct compartments: the dark zone (DZ), where antigen-activated B cells undergo proliferation and somatic hypermutation (SHM) to generate high-affinity B-cell receptors, and the light zone (LZ), where B cells undergo class-switch recombination (CSR) and are selected, based on antigen affinity, to differentiate into plasma cells or memory B cells^3^. DLBCL pathogenesis reflects the dysregulated accumulation of genetic alterations arising from repeated cycles of SHM, CSR, and proliferation, giving rise to two biologically distinct molecular subtypes: activated B-cell (ABC) and germinal center B-cell (GCB) DLBCL.

Consistent with global cancer trends, lymphoma was the fourth and fifth most common cancer among males and females, respectively, in Singapore between 2018 and 2022^4^, and the most common cancer diagnosis in males under 40 and females under 30.

### 1.2 Current treatment for DLBCL and limitations

First-line treatment for DLBCL is R-CHOP, a chemoimmunotherapy regimen combining rituximab, an anti-CD20 monoclonal antibody, with cyclophosphamide, doxorubicin, vincristine, and prednisolone. R-CHOP has remained the standard of care for two decades^5^. Despite substantially improving outcomes, 40–50% of patients develop intrinsic or acquired resistance to R-CHOP, making treatment failure the leading cause of morbidity and mortality in DLBCL^6^. Identifying biomarkers of treatment response and novel therapeutic strategies for relapsed/refractory (R/R) disease therefore remains an urgent need. However, despite advances in genomic profiling, the molecular mechanisms driving treatment failure are still poorly understood, limiting the development of effective targeted and combination therapies for R/R DLBCL.

Precision medicine and immunotherapy have transformed treatment across many cancer types in recent years, with immune checkpoint blockade emerging as a particularly promising strategy for overcoming immune evasion in prognostically adverse cancers.

### 1.3 Triple Negative Breast Cancer

Beyond hematological malignancies, dysregulated immune checkpoint signalling is similarly implicated in solid tumours such as triple-negative breast cancer (TNBC), a disease that, like DLBCL, faces high rates of treatment failure despite standard-of-care therapy. In Singapore, breast cancer accounts for both the highest incidence and highest mortality among females between 2018 and 2022^4^, consistent with global cancer statistics. TNBC, the most aggressive breast cancer subtype, is defined by the absence of estrogen receptor (ER), progesterone receptor (PR), and human epidermal growth factor receptor 2 (HER2) expression^7^. Its pathogenesis involves dysregulation of multiple signalling pathways, including PI3K/AKT/mTOR, Notch, MAPK/ERK, and JAK/STAT, which collectively drive tumour proliferation, progression, and metastasis^8^.

### 1.4 Current treatment for TNBC and limitations

TNBC accounts for approximately 40% of breast cancer-related mortality^9^, despite the availability of standard chemotherapy regimens built around taxanes, platinum agents, or anthracyclines as the cornerstone of breast cancer treatment^10^. As TNBC lacks ER/PR and HER2 expression, the targeted therapies effective in other breast cancer subtypes cannot be applied^11^. Many TNBC patients also develop chemoresistance or relapse despite an initial clinical response, contributing to the poorest prognosis of any breast cancer subtype. Immune checkpoint inhibitors targeting programmed death-ligand 1 (PD-L1) have improved outcomes in early-stage, non-metastatic TNBC^12^, but remain the only approved targeted therapy for this disease. Treatment options for advanced or metastatic TNBC therefore remain especially limited.

### 1.5 CD47 and its clinical significance

Cluster of differentiation 47 (CD47) is a ubiquitously expressed transmembrane protein of the immunoglobulin superfamily that functions as an innate immune checkpoint. By engaging signal regulatory protein alpha (SIRPα) on phagocytes, CD47 delivers an inhibitory “don’t-eat-me” signal that marks host cells as “self,” suppressing phagocytosis and protecting healthy autologous cells from inappropriate clearance^13^. Cancer cells exploit this mechanism: CD47 is overexpressed across both hematological and solid malignancies, including NHL, chronic myeloid leukemia, myeloma, osteosarcoma, and breast cancer^14^, where it engages SIRPα on myeloid cells to evade immunosurveillance. CD47 overexpression is broadly associated with poor clinical prognosis, including in therapy-resistant disease. In DLBCL specifically, CD47 is overexpressed relative to normal B cells and predicts poor prognosis, an effect attributed mainly to suppressed macrophage-mediated phagocytosis of malignant cells^15^. In TNBC, CD47 overexpression similarly correlates with metastasis and disease recurrence, and independently predicts poor survival^16^.

### 1.6 Non-immune functions of CD47: rationale for this study

Despite the well-established anti-phagocytic role of CD47, its downstream effects within tumour cells themselves, independent of immune interactions, remain poorly characterised. Emerging evidence suggests CD47 signalling also influences tumour-intrinsic pathways: CD47 has been implicated in mitochondrial biogenesis^17^ and glucose metabolism^18^, and low expression of the glycolytic enzyme glyceraldehyde-3-phosphate dehydrogenase (GAPDH) is prognostic for poor R-CHOP outcomes in DLBCL^19^, hinting at a link between CD47, metabolism, and treatment response. CD47 has also been shown to induce ferroptosis, an iron-dependent form of lipid peroxidation-driven cell death regulated by cellular metabolic state^20^, further suggesting that CD47 shapes tumour biology through metabolic and cell-intrinsic mechanisms extending beyond immune evasion. Given that CD47 is overexpressed and prognostically significant in both a haematological malignancy (DLBCL) and a solid tumour (TNBC), we hypothesised that CD47 also regulates non-immunological, pro-tumorigenic processes intrinsic to cancer cells, and that these effects may differ by cancer type and cellular context. In this study, we investigated the tumour-intrinsic consequences of CD47 loss in DLBCL and TNBC cell line models, characterising its effects on mitochondrial metabolism and chemosensitivity in DLBCL, and on cell cycle progression, migration, and drug response in TNBC. By defining these non-immune functions, we aim to expand current understanding of the cellular effects of CD47-targeted therapy and inform the rational design of combination treatment strategies.

## 2. Materials and Methods

### 2.1 Cell culture

Human DLBCL cell lines K231 and Toledo were cultured in Roswell Park Memorial Institute (RPMI) medium 1640. Human TNBC cell line MDA-MB-468 was cultured in Dulbecco’s Modified Eagle Medium/Nutrient Mixture F-12 (DMEM/F12) (Cytiva, USA). All media were supplemented with 10% fetal bovine serum (FBS) and 1% penicillin/streptomycin solution. All cells were cultured in a 5% CO_2_ humidified incubator at 37° Celsius.

### 2.2 Knockdown of CD47 via siRNA transfection

Cells were harvested and resuspended in fresh culture medium at 400,000 cells/mL. 200,000 cells/well were seeded onto a 24-well plate and transfected with 1 μL/well of control siRNA (cat. sc-37007, Santa Cruz Biotechnology, USA) or siRNA targeting CD47 (cat. sc-35006, Santa Cruz Biotechnology, USA) with Lipofectamine 3000 (cat no. L3000001, ThermoFisher Scientific, USA) and Opti-MEM according to the manufacturer’s protocol. Cells were then incubated overnight at 37°C, 5% CO_2_.

### 2.3 Analysis of CD47 expression via flow cytometry and fluorescence-activated cell sorting (FACS)

Flow cytometry was performed on transfected cells using a BD LSRFortessa X-20 Cytometer (BD Biosciences, USA) to validate CD47 silencing. Cells were stained with 0.2% R-PE (R-phycoerythrin) murine anti-human CD47 antibody in 10% FBS/PBS mixture (cat. 558046, BD Biosciences, USA) and incubated for 30 minutes at 4°C. Cells were then washed and resuspended in precooled phosphate-buffered saline (PBS) for flow cytometry. CD47-knockdown cells were isolated by FACS on the BD FACSVantage cell sorter (BD Biosciences, USA). Dead cells were excluded by staining for propidium iodide. Data were acquired using the FACSDiva 7.0 software (BD Biosciences, USA) and analysed using the FlowJo 10 software (BD Biosciences, USA). The experiments were performed in at least triplicates.

### 2.4 Analysis of cell cycle progression

Cells were harvested, washed with precooled PBS and fixed in 400 μL of 70% ethanol overnight at 4°C. The cells were then washed with PBS and resuspended in 400 μL of PBS containing 0.1% Triton X-100 (CAS No. 9036-19-5, Sigma Aldrich, USA), 20 μg/mL propidium iodide (CAS No. 25535-16-4, Sigma Aldrich, USA) and 0.2 mg/mL RNase A (cat no.10109142001, Sigma Aldrich, USA). Cells were incubated at 37°C for 15 minutes while protected from light before analysis by flow cytometry.

### 2.5 Oxygen consumption rate (OCR) and extracellular acidification rate (ECAR) assays

OCR and ECAR were measured on the Seahorse XFe96 Analyzer (Agilent Technologies, Santa Clara, CA, USA) using Seahorse XF Cell Mito Stress Test, according to the manufacturer’s instructions. Briefly, on the day of the experiment, 250,000 cells/well were seeded in 24-well Seahorse cell culture microplates, precoated with Corning™ Cell-Tak (Sacco, Cadorago, Italy). Cells were seeded in XF RPMI Medium (pH 7.4) supplemented with 1 mM pyruvate, 2 mM L-glutamine, and 10 mM glucose for OCR measurements, or with 2 mM L-glutamine for ECAR measurements, followed by the addition of medium to a final volume of 500 μL. The plates were incubated at 37 °C for one hour in a non-CO_2_ incubator. For OCR measurements, the Mito Stress Test was performed through sequential series of injections, starting with oligomycin (1 μM, ATP synthase inhibitor, port A), followed by carbonyl cyanide 4-(trifluoromethoxy) phenylhydrazone (FCCP, 2 μM, mitochondrial uncoupler, port B), and a combination of antimycin and rotenone (0.5 μM, CIII and CI inhibitors, port C). For ECAR measurements, the glycolytic stress test was performed via injection of glucose (10 mM, glycolysis substrate, port A), followed by oligomycin (1 μM, port B), and 2-deoxy-D-glucose (2DG, 50 mM, glycolysis inhibitor, port C). OCR and ECAR readings were recorded after every injection. Respiratory parameters were calculated with the Wave Desktop 2.6 software and shown as mean ± SEM of three independent experiments.

### 2.6 Colony formation assay

300 cells were seeded onto a 6-well cell culture dish and incubated at 37°C for 7 days. The wells were first washed with PBS, colonies were fixed with 10% neutral buffered formalin solution for 15 minutes and then stained with 1% crystal violet solution (Sigma, USA) for 30 minutes. Wells were thoroughly washed with distilled water and allowed to dry overnight. Digital images of the colonies were captured using a brightfield microscope.

### 2.7 Macrophage phagocytosis assay

U937 cells were seeded at a density of 15,000 cells/well in a 96-well microtiter plate. The cells were allowed to be differentiated into macrophages following exposure to 100 nM phorbol 12-myristate 13-acetate (PMA) (Sigma, USA) for 5 days. Macrophages were then labelled with carboxyfluorescein succinimidyl ester (CFSE) dye (Thermo Fisher Scientific, USA) while CD47 siRNA-transfected DLBCL cells were labelled with pH-sensitive dye, pHrodo Red (which emitted a red fluorescence upon a drop in pH when engulfed cells were inside phagosomes) (Thermo Fisher Scientific, USA). Subsequently, DLBCL cells were incubated with either IgG1 (10 μg/mL, Santa Cruz Biotechnology, USA) or anti-CD47 antibody (10 μg/mL, B6H12 clone, Santa Cruz Biotechnology, USA) for one hour at room temperature. Pre-seeded macrophages and labelled DLBCL cells were combined at a 1:5 effector-to-target ratio, before incubation at 37°C for two hours. The wells were washed twice with PBS to remove non-adherent and non-phagocytosed DLBCL cells. BX61 fluorescent microscope (Olympus, Japan) was used to assess phagocytosis.

The following formulas were utilised:

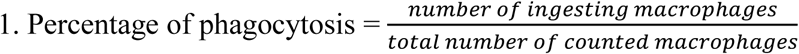

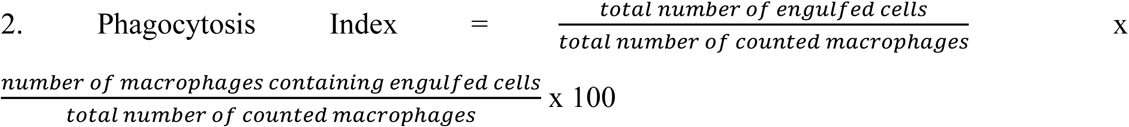

Five random fields of view were assessed for quantification of each condition. Two experiments were conducted with each condition analysed in duplicates.

### 2.8 Drug response assay

Drug response assays were performed to compare the sensitivity of cells to doxorubicin (doxorubicin hydrochloride 10 mg, cat no. 25316-40-9, Sigma Aldrich, USA), paclitaxel (Abraxane® 100 mg/vial), docetaxel (Hospira^TM^, 20 mg/2ml) and oxaliplatin (Hospira^TM^, 5 mg/ml) for TNBC cells and rituximab, doxorubicin, cyclophosphamide, prednisone and vincristine (R-CHOP) for DLBCL cells. A range of 10 concentrations of each drug was prepared via serial dilution in culture medium. Both control siRNA and CD47 siRNA-treated cells were seeded on a 96-well plate at 10,000 cells/well in 100 μL of medium/well and incubated for 24 hours at 37°C, 5% CO_2_. The cells were then treated with respective concentrations of drugs and incubated for another 48 hours. The respective cell viabilities were obtained using 100 μL/well of CellTiter-Glo 2.0 reagent according to the manufacturer’s protocol. The luminescent signal was recorded with Tecan Infinite 200 Pro microplate reader using an integration time of one second/well. Respective half-maximal inhibitory concentration (IC_50_) curves were plotted using GraphPad Prism 8.0.2 (GraphPad, USA).

### 2.9 Cell viability assay

10,000 cells transfected with either control siRNA or CD47 siRNA were suspended in 100 μL of fresh medium and seeded into each well in a NUNC flat-bottom white 96-well microplate (ThermoFisher, USA). Cells were then incubated for 48 hours at 37°C, 5% CO_2_. 100 μL/well of CellTiter-Glo 2.0 reagent (Promega, USA) was then used according to the manufacturer’s protocol. The luminescent signal was recorded with Tecan Infinite 200 Pro microplate reader using an integration time of one second/well.

### 2.10 Wound healing assay

100,000 cells were seeded on both sides of silicone inserts (Ibidi, Germany) adhered onto the wells of a 12-well plate and incubated for 48 hours at 37°C, 5% CO_2_ to produce confluent monolayers separated by a 500 µm gap in each well. Inserts were removed and gaps were photographed using light microscopy and further incubated for 48 hours. Images of the gap were collected at the 0-hour, 24-hour and 48-hour marks. The migration rate was determined with ImageJ software using the uncovered surface area of the gaps.

### 2.11 Western blotting

Whole cell lysates were obtained by lysing cells in radioimmunoprecipitation lysis and extraction buffer (cat. 89900, ThermoFisher Scientific, USA) supplemented with 1x protease inhibitor cocktail (cat. 78430, ThermoFisher Scientific, USA). Protein concentrations of cell lysates were quantified using Bradford protein assay (cat. 5000201, Bio-Rad, USA) according to the manufacturer’s protocol. Lysate containing 20 μg of protein from each sample was subjected to electrophoretic fractionation in 10% SDS-polyacrylamide gel before transferring to polyvinylidene difluoride membranes (Bio-Rad, USA) using the Trans-Blot® Turbo^TM^ Transfer System (Bio-Rad, USA). After incubation with 5% non-fat milk in Tris-buffered saline with Tween 20 (TBST) (10mM Tris, pH 8.0, 150mM NaCl, 0.5% Tween 20) for 60 minutes, the membrane was washed once with TBST and incubated with murine anti-CD47 antibody B6H12 (cat. sc-12730, Santa Cruz Biotechnology, USA) or beta-actin antibody at 4°C overnight. Following overnight incubation, the membranes were washed five times with TBST and incubated with horseradish peroxidase-conjugated anti-mouse secondary antibodies (cat. sc-358914, Santa Cruz Biotechnology, USA) at a dilution factor of 1:2000 for two hours. The membranes were then washed five times with TBST and once with TBS, before visualising the bands with enhanced chemiluminescence detection reagents (ThermoFisher Scientific, USA) and detecting with an iBright^TM^ FL1500 Imaging System (ThermoFisher Scientific, USA).

### 2.12 RNA sequencing

Total RNA was extracted using the RNEasy Plus Mini Kit (Qiagen, USA). At least 1000ng of RNA was used for the subsequent workflow. RNA libraries were prepared using the NEBNext® Ultra™ II Directional RNA Library Prep Kit for Illumina (New England BioLabs, USA). Sequencing was performed on the NovaSeq 6000 platform (Illumina, USA). Library preparation, sequencing and analysis were carried out at Novogene (Singapore).

### 2.13 Immunohistochemistry (IHC) staining and CD47 quantification

IHC staining was performed on formalin-fixed paraffin-embedded (FFPE) tissue samples using anti-CD47 monoclonal antibody (cat. ab226837, clone SP279, Abcam, UK) on the Leica Bond autostainer at the NUH Department of Pathology. Antigen retrieval was performed using pH 9 for 20 minutes and the samples were incubated for 15 minutes with the primary antibody at a dilution of 1:50, according to the manufacturer’s protocol. Quantification of CD47 protein expression by 3,3′-diaminobenzidine (DAB) staining was conducted on QuPath v0.3.

### 2.14 Knockout of CD47 via CRISPR-Cas9 system

CRISPR knockout (cat. sc-400508-KO-2) and control CRISPR (cat. sc-418922) lentiviral constructs (Santa Cruz Biotechnologies, USA) were transfected into HEK293FT cells to produce lentiviruses. Lentiviruses were then concentrated using Lenti-X concentrator (Takara Bio, USA) before being added to the cell lines, supplemented with 8 µg/mL polybrene (Sigma Aldrich, USA).

### 2.15 Statistical analysis

Statistical analyses were performed using GraphPad Prism 8.0.2. Normality of data was evaluated using the Shapiro-Wilk test. Normally distributed data were analysed using the two-tailed independent samples t-test. A p-value ≤ 0.05 was considered statistically significant.

## 3. Results

### 3.1 CD47 in DLBCL

#### 3.1.1 CD47 expression is correlated with clinical outcomes and subtypes in DLBCL patients

Using publicly available data from The Cancer Genome Atlas (TCGA), DLBCL tumours were found to express higher levels of CD47 than normal B cells (Figure 1A), and high CD47 gene expression correlated with poorer overall survival (Figure 1B). We then characterised CD47 protein and mRNA expression by IHC and RNAscope in-situ hybridisation (ISH), respectively, in de novo DLBCL patient FFPE samples from NUH, constructed as a tissue microarray (TMA). Consistent with the TCGA data, patients with high CD47 mRNA expression showed significantly worse progression-free survival (PFS) and overall survival (OS) (Figure 1C), and DLBCL patients who relapsed after R-CHOP treatment expressed significantly higher CD47 mRNA levels (1.97× higher, p = 0.038) (Figure 1D). These correlations were not statistically significant for CD47 protein expression, a disparity that may point to a regulatory mechanism specific to the transcriptional stage of CD47. ABC-subtype DLBCL cases also expressed significantly higher CD47 levels than GCB-subtype cases [2.8× higher, p < 0.0001 (protein) and 2.38× higher, p = 0.0005 (RNA)] (Figure 2), suggesting subtype-specific CD47 signalling consistent with the poorer survival outcomes reported for ABC-subtype DLBCL. Since overexpression of both MYC and BCL2 are established prognostic markers of adverse outcomes in DLBCL, we stratified the NUH cohort by MYC and BCL2 positivity (via IHC) and examined the corresponding CD47 expression: CD47 mRNA levels were significantly higher in patients positive for MYC (p = 0.0285) or BCL2 (p = 0.0105) (Figure 3), pointing to a possible link between CD47 and these markers. Together, the consistent association between CD47 expression and poor survival across these independent analyses indicates a general dependence on CD47 for disease progression in DLBCL.

**Figure 1.**
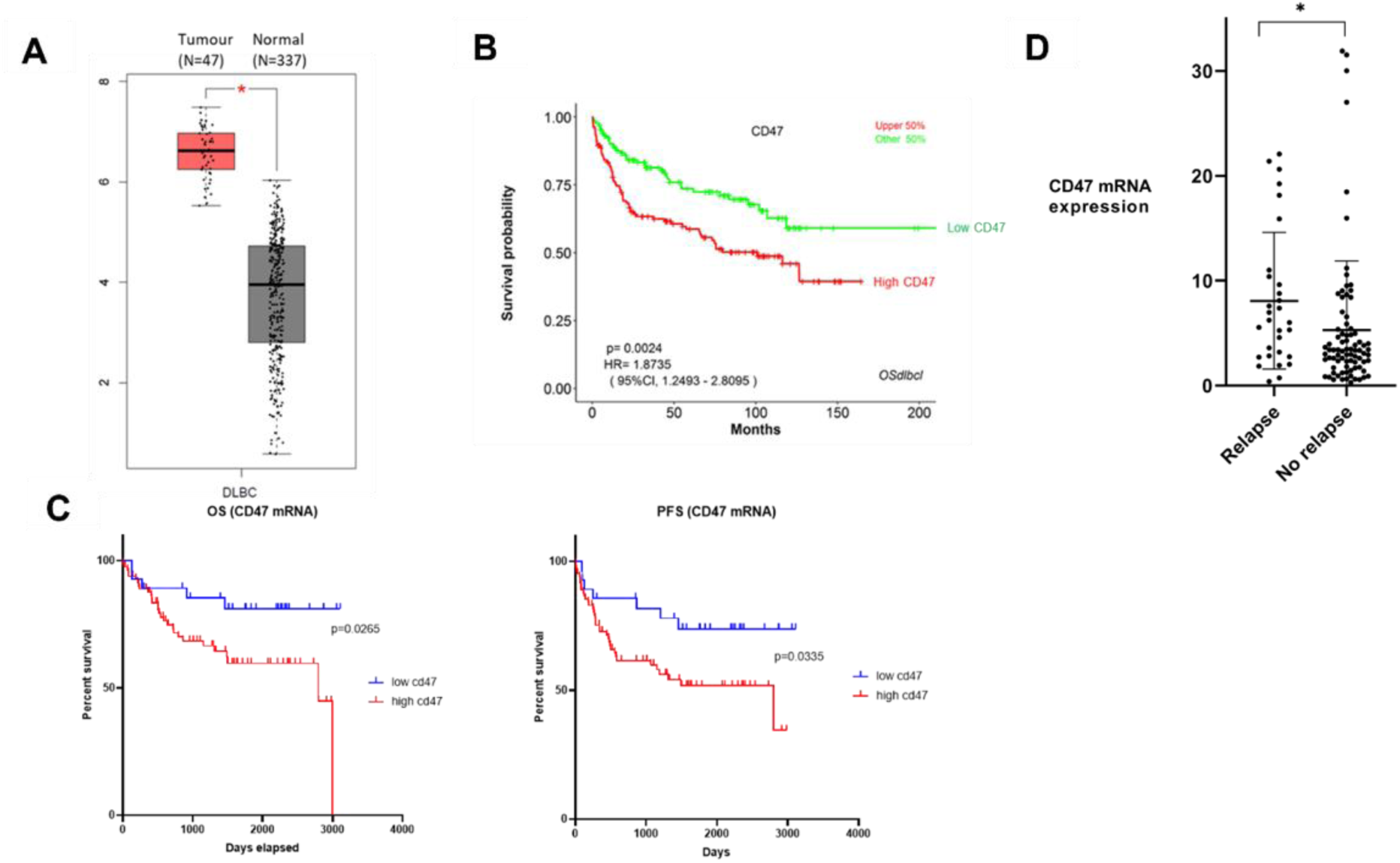
CD47 expression and corresponding survival in DLBCL patients. (A) CD47 gene expression in DLBCL tumours (n=47) versus normal B cells (n=337) and (B) overall survival curve from the TCGA dataset. (C) OS (left) and PFS (right) of DLBCL patients with high or low CD47 mRNA expression. (D) CD47 expression in DLBCL tumours from relapsed and non-relapsed patients, as quantified by RNAscope ISH assay applied to the NUH cohort.

**Figure 2.**
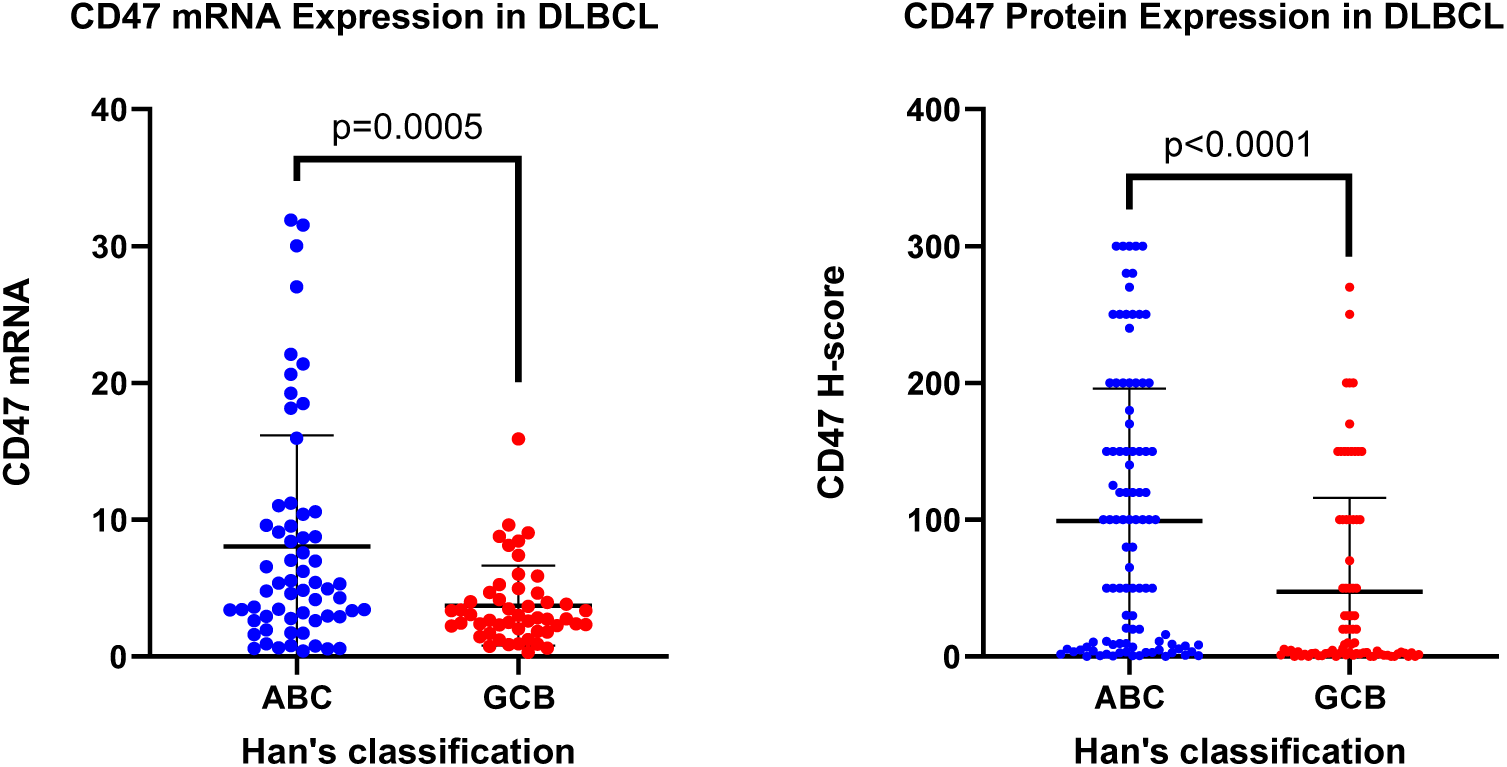
CD47 expression classified according to DLBCL subtype, as quantified by RNAscope for mRNA and by H-score for surface protein expression.

**Figure 3.**
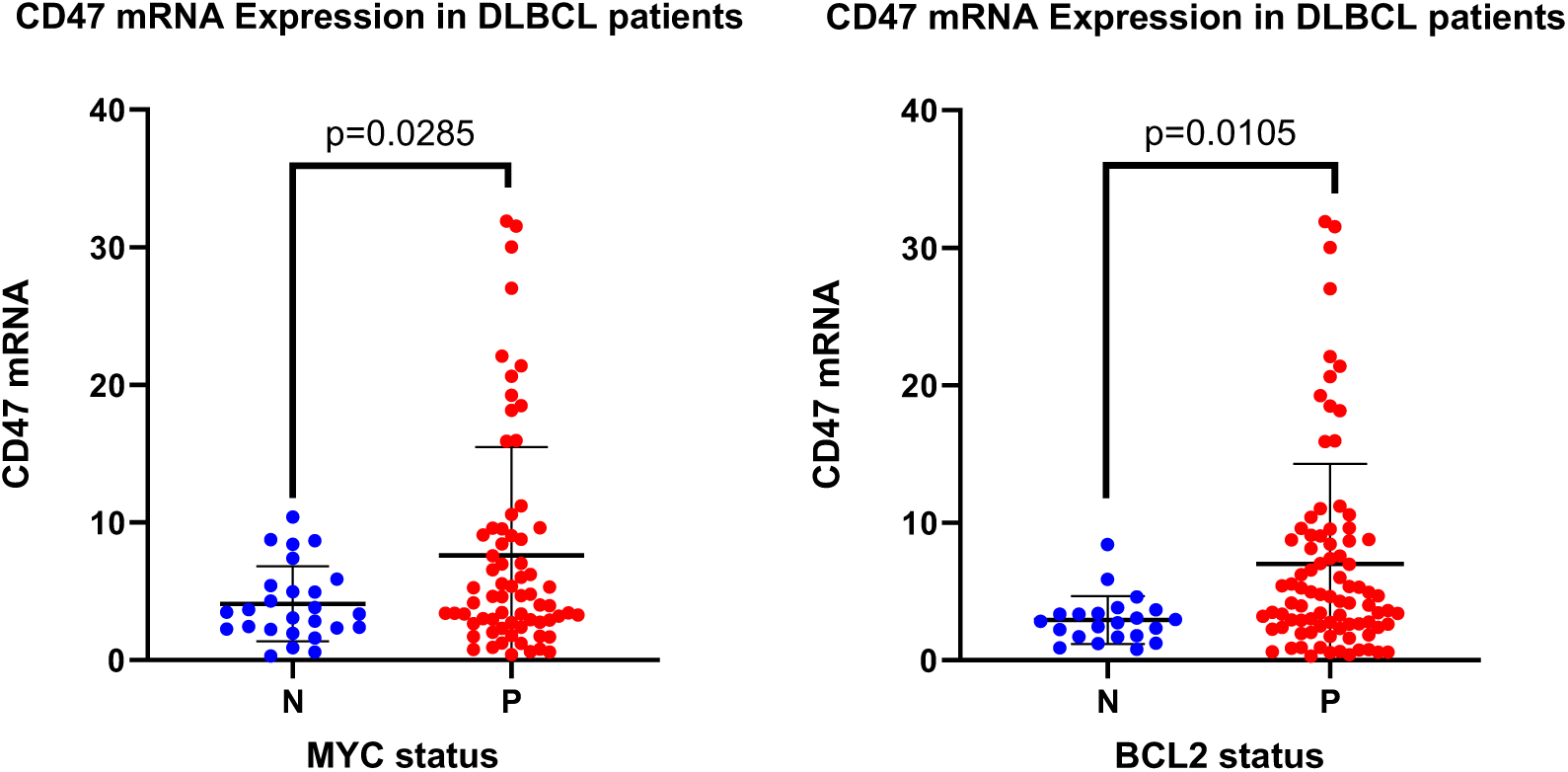
CD47 mRNA expression in patients from the NUH cohort stratified based on positive expression of MYC and BCL2 by IHC.

#### 3.1.2 CD47 mRNA and protein expressions were characterised in DLBCL cell lines

We next assessed CD47 expression across a panel of DLBCL cell lines to identify suitable models for generating CD47-modulated isogenic lines. CD47 mRNA and protein expression, determined by RNAscope ISH and western immunoblotting respectively, varied substantially across cell lines, mirroring the heterogeneity observed in patient samples (Figure 4). Based on their high CD47 expression, we selected K231 and Toledo to generate stable CD47 knockdown (via siRNA transfection) and knockout (via CRISPR-Cas9) models for downstream functional studies. Figure 4C shows CD47 knockout in the K231 line, generated by CRISPR-Cas9 lentiviral transduction and confirmed by flow cytometry.

**Figure 4.**
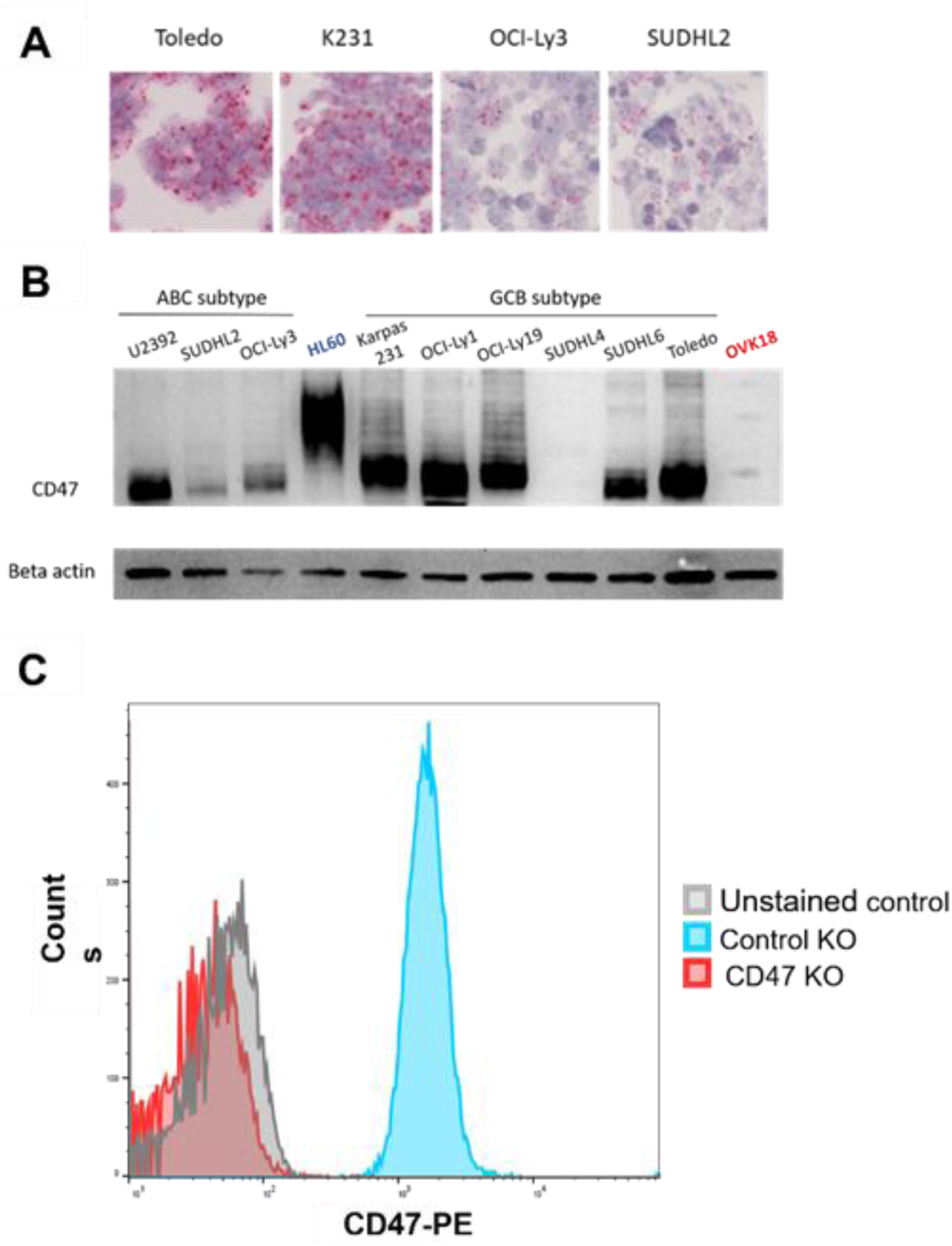
CD47 expression in DLBCL cell lines. (A) Representative RNAscope images of CD47 mRNA in highly expressing cell lines (Toledo and K231) and low expressors (OCI-Ly3 and SUDHL2). Each mRNA molecule is denoted by a pink dot, with hematoxylin staining for the nuclei, allowing for absolute quantification of mRNA per cell. (B) Western immunoblotting of representative DLBCL cancer cells across the ABC and GCB subtypes. The thick bands reflect varying extents of glycosylation of CD47. (C) Flow cytometry histograms depicting the generation of CD47 CRISPR knockout K231 cell line.

#### 3.1.3 CD47 regulates the metabolism of DLBCL cells

To identify pathways regulated by cell-intrinsic CD47 signalling, we performed RNA sequencing comparing control siRNA-treated and CD47 siRNA-treated K231 DLBCL cells, followed by pathway enrichment analysis (PEA) of the differentially expressed genes. The most significantly enriched pathways following CD47 knockdown were related to oxidative phosphorylation and mitochondrial function (Figure 5), suggesting that CD47 normally suppresses these metabolic pathways, potentially favouring a shift toward glycolytic dependence.

**Figure 5.**
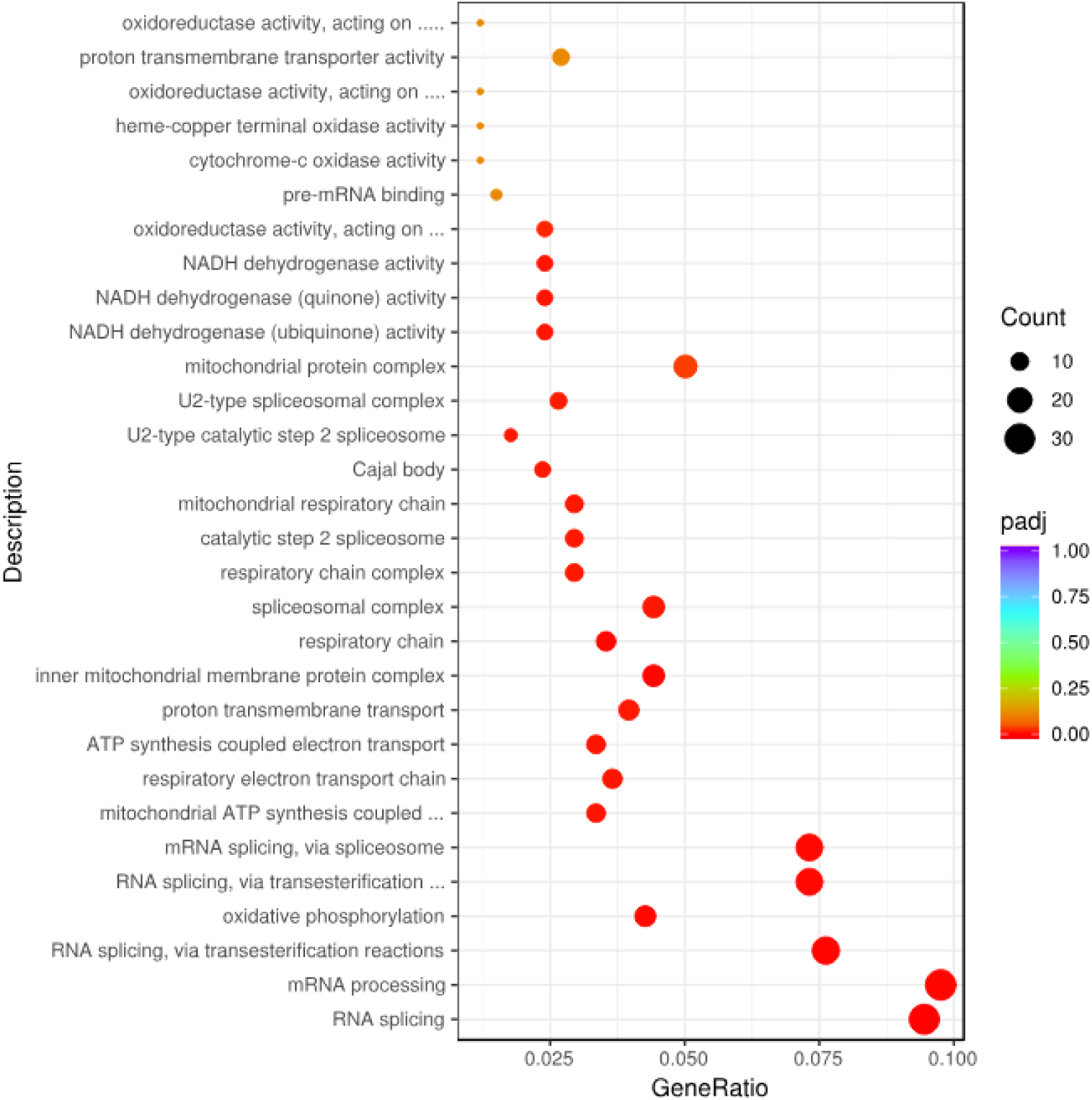
PEA analysis was performed on the differentially expressed genes. Common hits associated with loss of CD47 expression include oxidative phosphorylation and mitochondria-related pathways including respiratory electron transport.

#### 3.1.4 CD47 knockout reduced oxygen consumption rate (OCR) in DLBCL cell lines

To functionally validate the PEA findings, we assessed the effect of CD47 knockout on DLBCL cell metabolism using the Seahorse MitoStress assay, which measures oxygen consumption rate (OCR) and extracellular acidification rate (ECAR) alongside sequential mitochondrial inhibitor treatment. CD47 knockout consistently reduced OCR in both K231 and Toledo clones relative to their controls (Figure 6), directly linking CD47 to basal mitochondrial respiration and indicating mitochondrial dysfunction with reduced oxidative ATP output. ECAR was unchanged in both cell lines, indicating that glycolytic activity was unaffected by CD47 loss.

**Figure 6.**
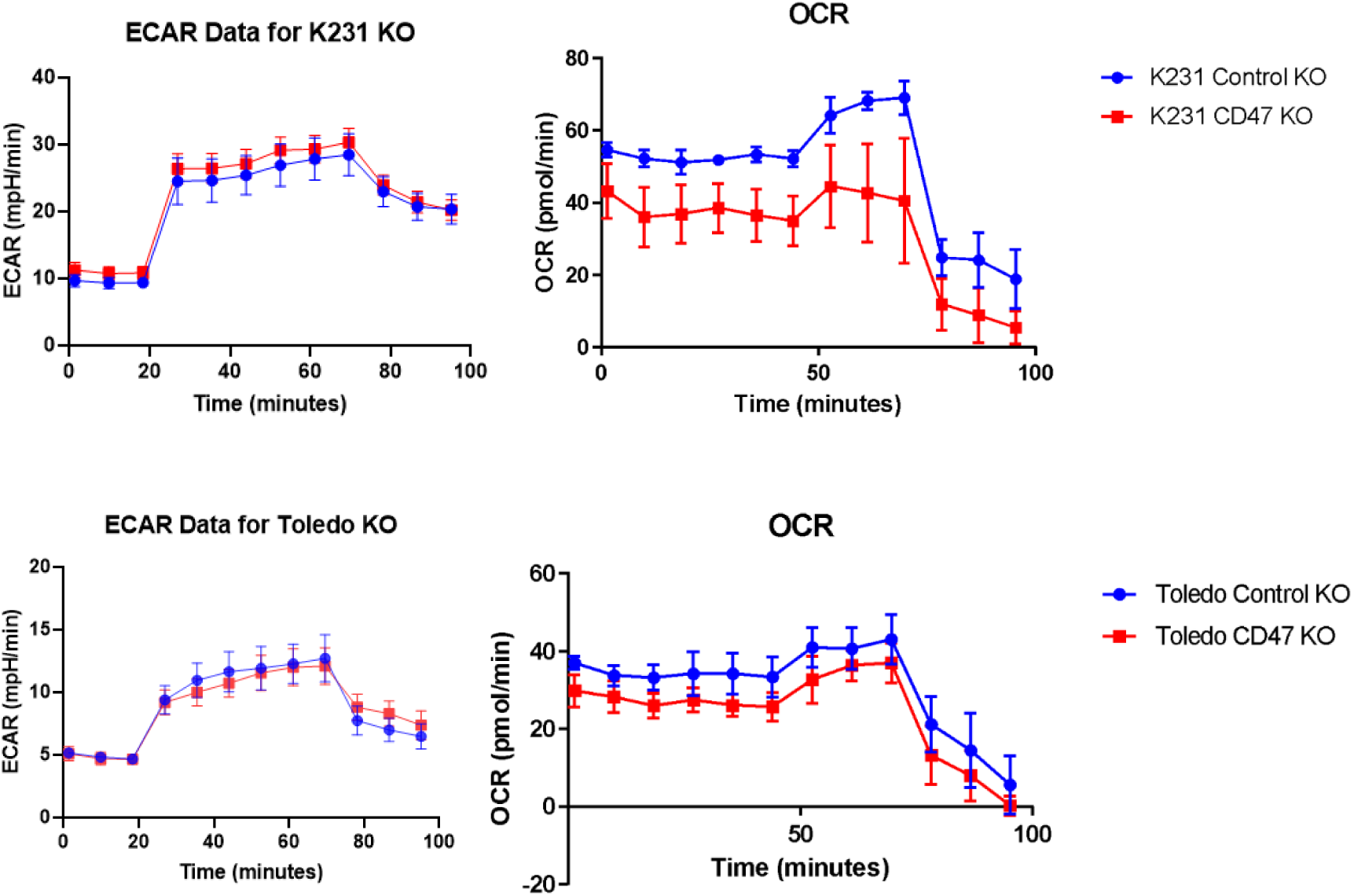
Effect of CD47 knockout on ECAR and OCR in K231 (top) and Toledo (bottom) cells, as determined by Seahorse MitoStress assay.

#### 3.1.5 Reduced CD47 expression in DLBCL cells increased sensitivity towards R-CHOP therapy

We next investigated whether CD47 expression influences DLBCL cell sensitivity to R-CHOP, the first-line chemo-immunotherapy used clinically for DLBCL. Dose-response curves showed a lower R-CHOP IC50 in CD47 knockout K231 cells relative to controls, but only in the presence of human serum. No significant change was observed in the presence of FBS (Figure 7).

**Figure 7.**
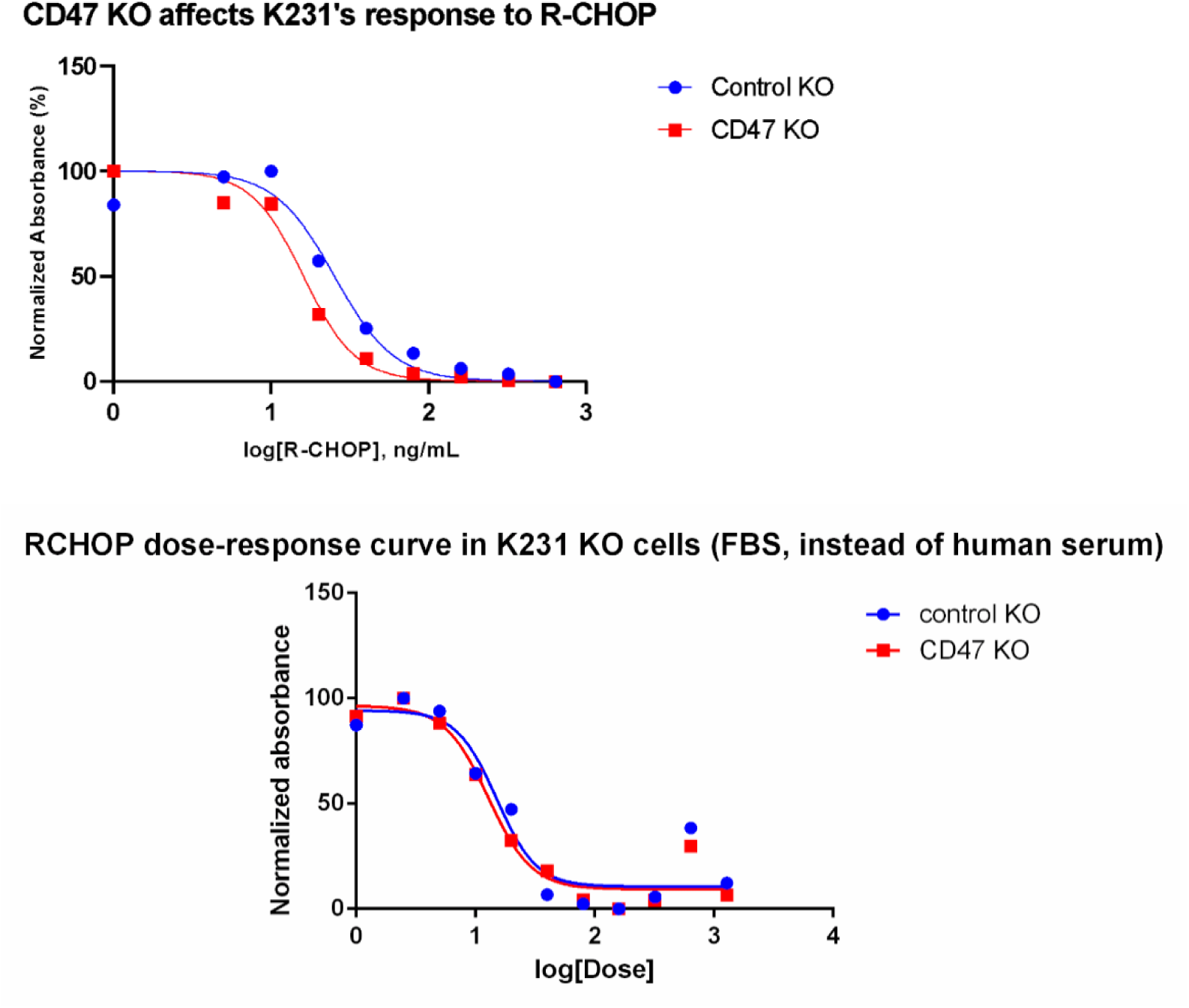
Dose-response relationships of viability of control knockout versus CD47 knockout K231 cells upon R-CHOP treatment as determined via drug-response assays. Dose response curves of control knockout and CD47 knockout K231 cells upon R-CHOP therapy in the presence of human serum (top) and FBS (bottom).

#### 3.1.6 Reduced CD47 expression in DLBCL cells increased dysfunctional mitochondria

The JC-1 assay measures mitochondrial membrane potential by flow cytometry: in healthy cells, JC-1 forms aggregates within mitochondria that fluoresce red, whereas in unhealthy cells, JC-1 monomers remain in the cytoplasm and fluoresce green (Figure 8A). At baseline, without R-CHOP treatment, CD47 knockout DLBCL cells showed a higher proportion of unhealthy mitochondria than controls (Figure 8B), indicating that basal mitochondrial health depends on CD47.

**Figure 8.**
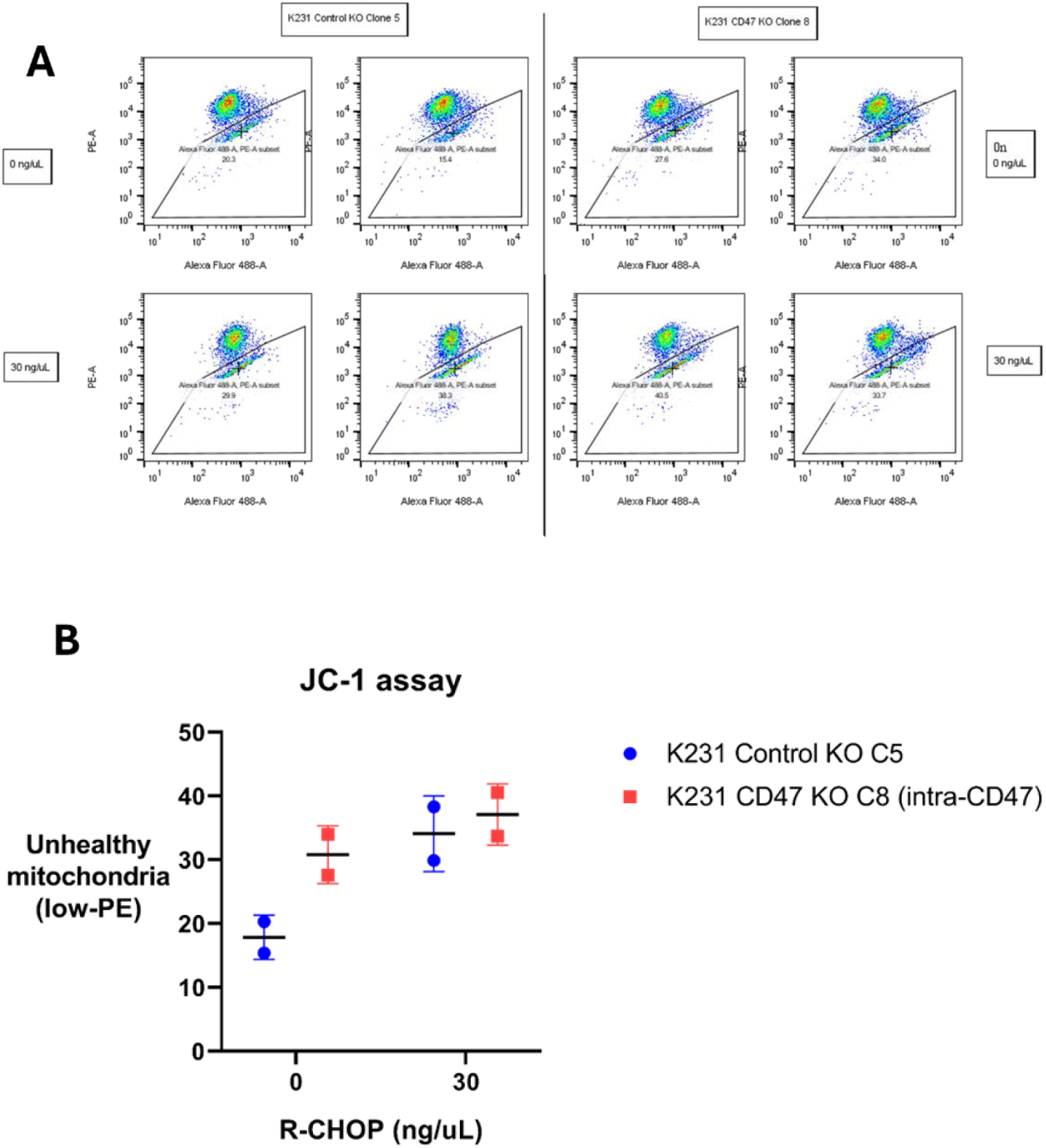
JC-1 assay was performed to assess the proportion of dysfunctional mitochondria. (A) A gating strategy was first established on the flow cytometer. (B) The assay was then performed on K231 cells with and without R-CHOP treatment.

#### 3.1.7 CD47 knockout increased apoptotic rate in DLBCL cells upon R-CHOP treatment

CD47 knockout K231 cells showed higher levels of early apoptosis following treatment with 30 ng/μL R-CHOP, indicating increased sensitivity to the chemo-immunotherapy (Figure 9).

**Figure 9.**
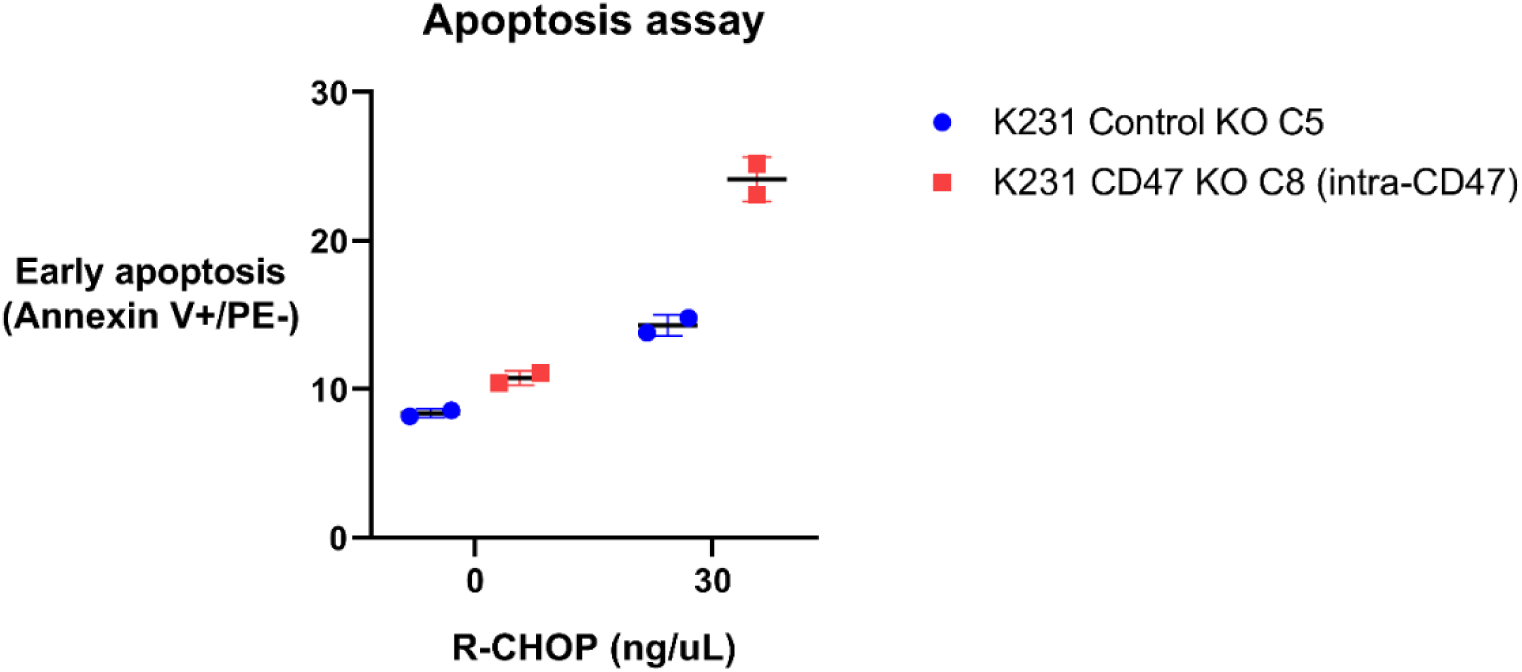
Apoptosis assay was performed to compare apoptosis in K231 cells with and without R-CHOP therapy.

#### 3.1.8 Western blot analysis confirmed downregulation of mitochondria-related and other genes in CD47 knockout DLBCL cells

Immunoblotting of candidate genes identified from the RNA-sequencing data confirmed downregulation of GAS8 (involved in the Hedgehog signalling cascade) and ABCB1 following CD47 loss (Figure 10). We also examined the mitochondria-related genes PGC1α (master regulator of mitochondrial biogenesis) and DRP1 (regulator of mitochondrial fission), and found an unexpected, paradoxical increase in full-length PGC1α with no change in DRP1 (Figure 10), suggesting that CD47 downregulation affects mitochondrial activity rather than mitochondrial gene expression per se.

**Figure 10.**
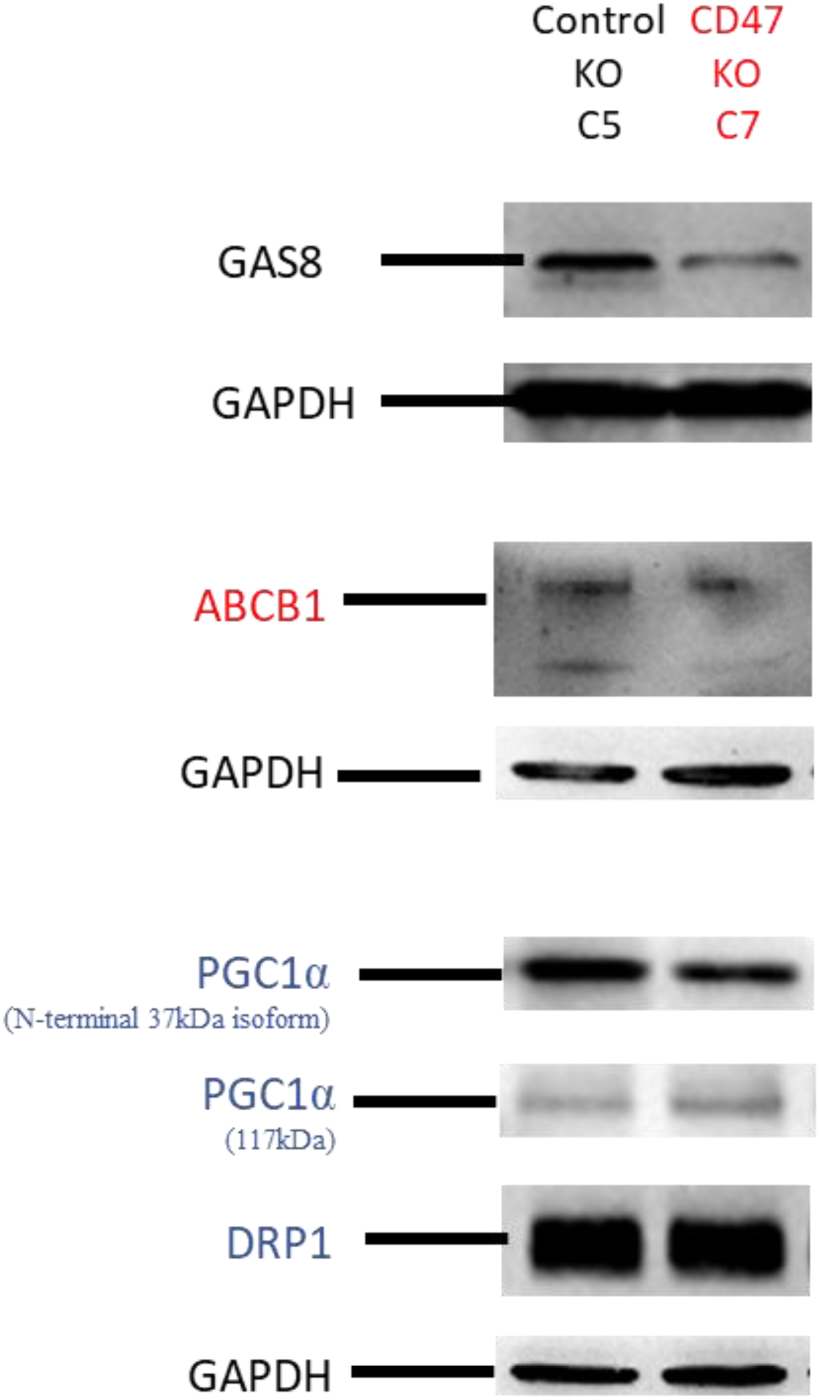
Expression of candidate genes from RNA-sequencing data and mitochondria-related genes in control or CD47 knockout K231 cells as determined by Western blot analysis.

#### 3.1.9 Western blot analysis indicated possible involvement of CD47 in Hedgehog signalling pathway in DLBCL

Given the downregulation of GAS8, we examined its downstream effect on the Hedgehog signalling cascade in K231 cells. Immunoblotting of Hedgehog pathway proteins showed that CD47 knockout downregulated Shh, the penultimate ligand of the cascade (Figure 11).

**Figure 11.**
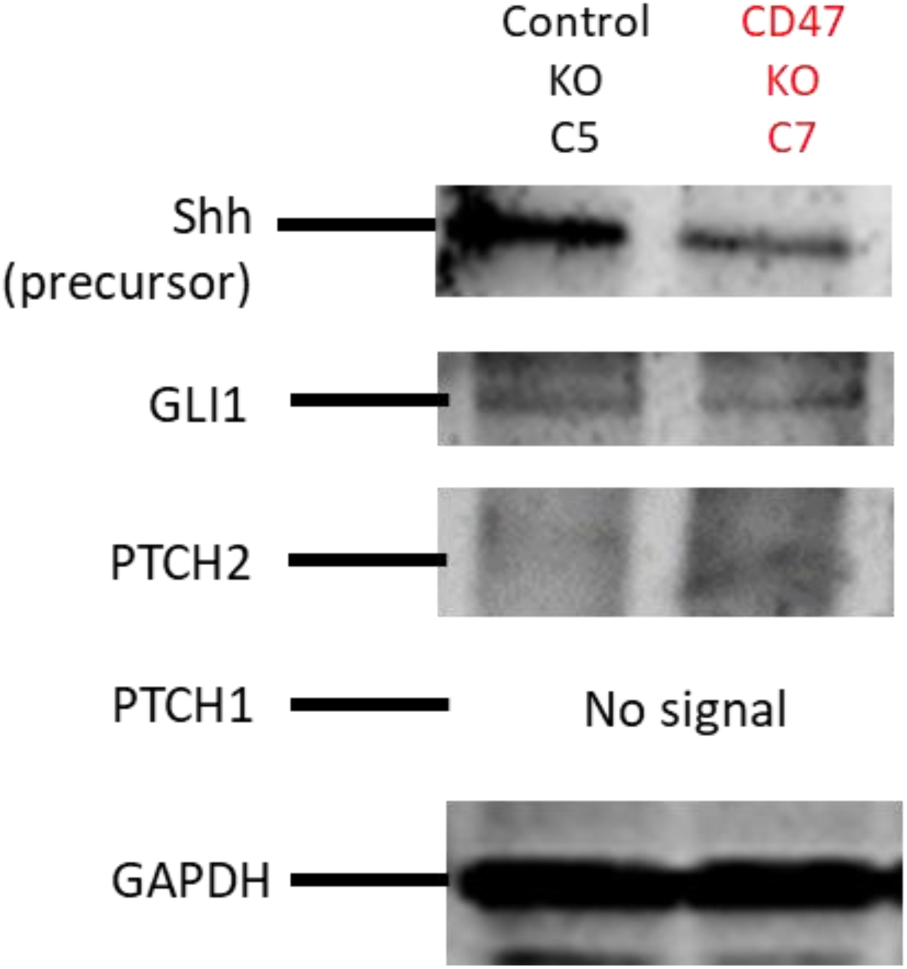
Expression of Hedgehog signalling pathway proteins in control or CD47 knockout K231 cells as determined by Western blot analysis.

### 3.2 CD47 in TNBC

#### 3.2.1 CD47 expression was intrinsically upregulated in MDA-MB-468 cells

As MDA-MB-468 is a widely used TNBC cell line model, we first confirmed that it expresses higher CD47 than MCF-10A, a normal immortalised breast epithelial line. Both are human mammary cells of epithelial morphology, allowing direct comparison of CD47 expression in TNBC versus normal breast tissue. Flow cytometry showed CD47 expression in MDA-MB-468 cells was 45.1 ± 2.34-fold higher than in MCF-10A (Figure 12), consistent with previously reported elevated CD47 expression in TNBC. Having confirmed high CD47 expression in MDA-MB-468 cells, we proceeded to CD47 knockdown experiments to determine its effects on cellular activity.

**Figure 12.** CD47 expression in MDA-MB-468 and MCF-10A cells as determined by flow cytometric analysis. (A) Representative flow cytometric histogram plots showing CD47 expression on normal immortalised breast cells MCF-10A (red) and TNBC MDA-MB-468 cells (blue). (B) Bar graph depicting percentage of fluorescence intensity of surface CD47 in MDA-MB-468 and MCF-10A cells, normalised to the mean fluorescence intensity of CD47 expression in MDA-MB-468 cells. Data represents mean ± S.E.M. from three independent experiments (n=3). ****p < 0.0001.

#### 3.2.2 siRNA transfection reduced CD47 expression in MDA-MB-468 cells

To confirm siRNA-mediated CD47 knockdown, we compared cells transfected with CD47 siRNA against control siRNA. Cell-surface CD47 was assessed by flow cytometry and immunocytochemical staining, and total CD47 protein by western immunoblotting (with GAPDH as a loading control). CD47 expression was significantly reduced in CD47 siRNA-transfected cells across all three methods (Figure 13).

**Figure 13.**
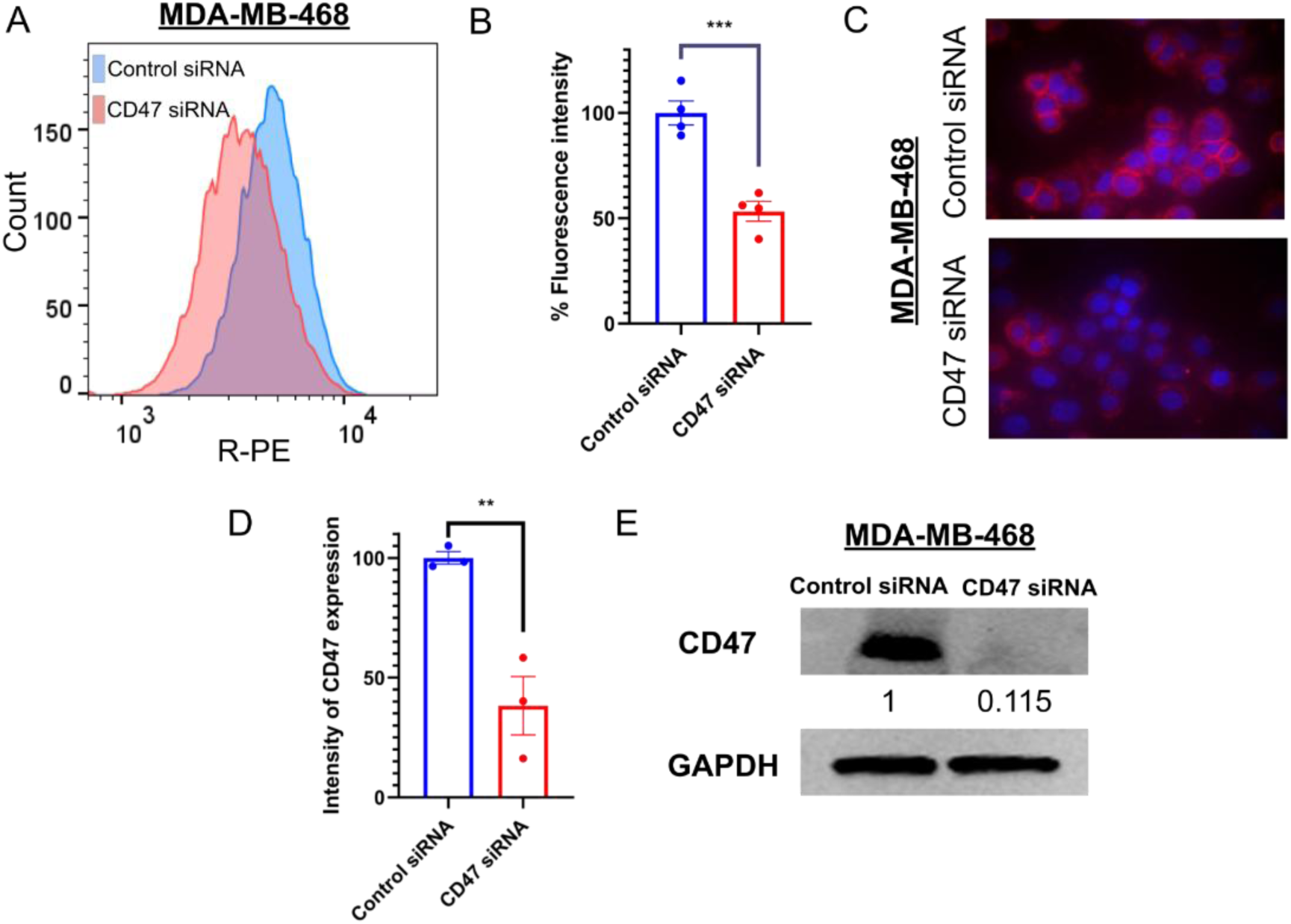
CD47 expression in MDA-MB-468 cells transfected with either control siRNA or CD47 siRNA as determined by flow cytometry, immunocytochemical staining and Western immunoblotting. (A) Representative flow cytometric histogram plots showing CD47 expression in MDA-MB-468 cells. Blue histogram represents MDA-MB-468 cells transfected with 1 μL of control siRNA with no homology to any eukaryotic gene. Red histogram represents MDA-MB-468 cells transfected with 1 μL of siRNA targeting the CD47 sequence. (B) Bar graph depicting percentage of fluorescence intensity of surface CD47 in MDA-MB-468 cells transfected with control siRNA (blue) and CD47 siRNA (red) normalised to intensities obtained for control siRNA-treated cells. Data represents mean ± S.E.M. of four independent experiments (n=4). (C) Immunocytochemical staining for surface CD47 (in red) in control siRNA-treated MDA-MB-468 cells (top) and CD47 siRNA-treated MDA-MB-468 cells (bottom). Cells were incubated with 0.2% R-PE mouse anti-human CD47 antibody. Nuclei (in blue) were counterstained with Hoechst 33342. (D) Bar graph quantification of fluorescence intensity depicting levels of CD47 expression for MDA-MB-468 cells transfected with either control or CD47 siRNA normalised to intensities obtained for control siRNA-treated cells. Data represents mean ± S.E.M. of three independent experiments (n=3). (E) Western blot analysis showing CD47 protein expression for control siRNA-treated MDA-MB-468 and CD47 siRNA-treated MDA-MB-468 cells. Housekeeping GAPDH was used as loading control. ***p < 0.001, **p < 0.01

#### 3.2.3 CD47 knockdown delayed cell cycle progression in MDA-MB-468 cells

Following siRNA transfection, a larger proportion of CD47 siRNA-treated cells appeared enlarged and flattened relative to controls (Figure 14A). Cell cycle analysis confirmed a significantly enriched G1 fraction in CD47 siRNA-treated cells (61.9 ± 2.96% versus 54.8 ± 0.686% in controls, p < 0.05) (Figure 14B, 14C). RNA sequencing also revealed significant downregulation of CCND1, which encodes cyclin D1, a key regulator that promotes the G1-to-S phase transition, a plausible driver of the observed cell cycle arrest.

**Figure 14.**
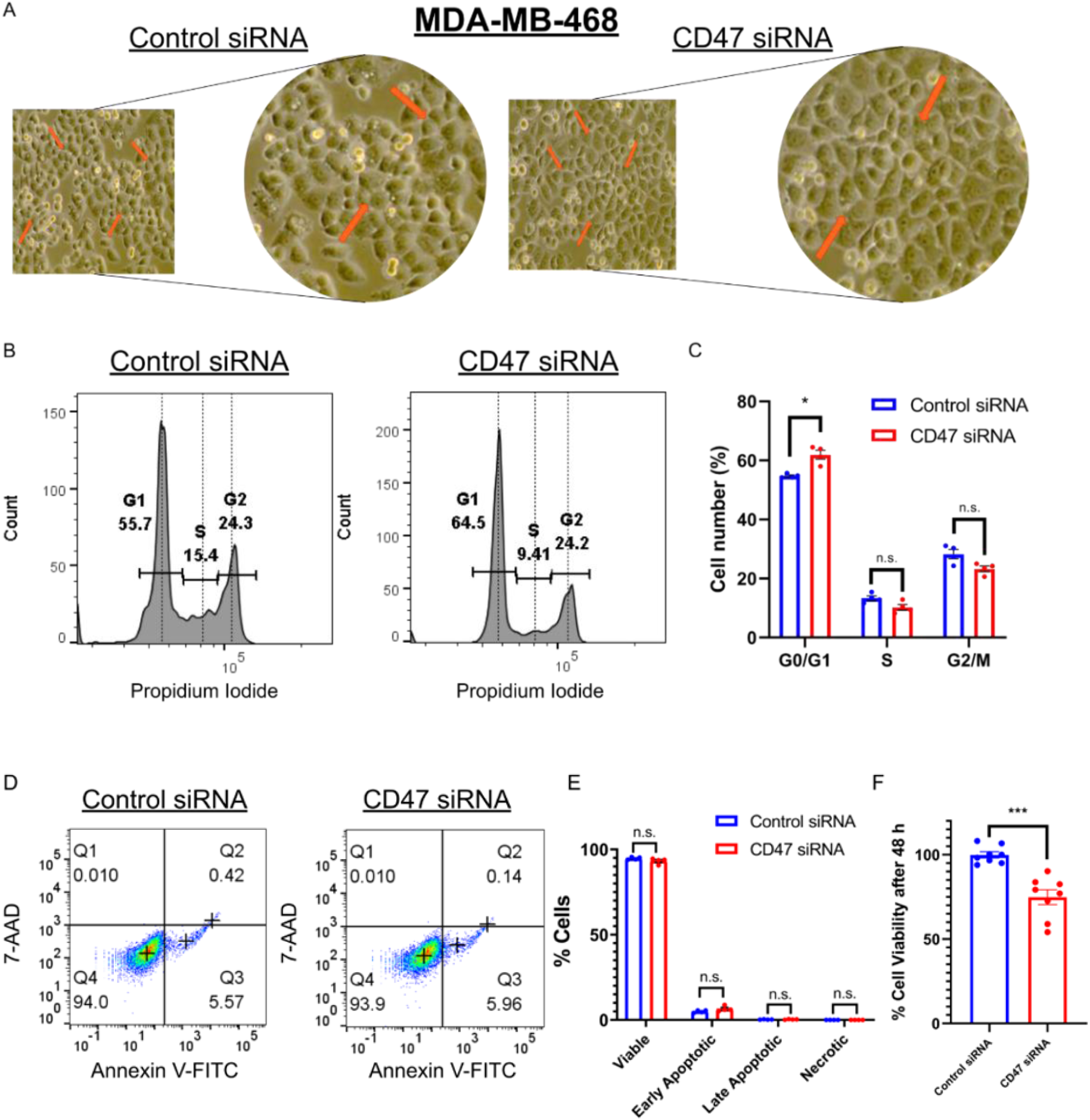
Effect of CD47 silencing on cell cycle, apoptosis and cell viability in MDA-MB-468 cells as determined by flow cytometric analysis and cell viability assay. (A) Representative light microscopic images of control siRNA-treated (left) and CD47 siRNA-treated MDA-MB-468 cells (right) obtained at 40x magnification. (B) Representative histogram plots for cell cycle analysis by flow cytometry of control siRNA-treated (left) and CD47 siRNA-treated MDA-MB-468 cells (right). (C) Bar graph representation of CD47 siRNA treatment having induced significant G0/G1 phase cell cycle arrest in MDA-MB-468 cells. Data represents the mean ± S.E.M. of four independent experiments (n=4). (D) Representative flow cytometric plots for apoptosis assay for control siRNA-treated (left) and CD47 siRNA-treated MDA-MB-468 cells (right). Cells were stained with 5 μL of Annexin V-fluorescein isothiocyanate (FITC) conjugate solution and 5 μL of 7-aminoactinomycin D (7-AAD). Average percentage of late and early apoptotic cells were reported in quadrant 2 and 3 respectively. (E) Bar graph representation of apoptosis results. Data represents the mean ± S.E.M. of four independent experiments (n=4). (F) Cell viability of control siRNA-treated (blue) and CD47 siRNA-treated MDA-MB-468 cells (red) were determined using the Cell-Titre-Glo Luminescent Assay 48 h after siRNA transfection. CellTiter-Glo 2.0 Reagent was added in a 1:1 ratio to the cell suspension and luminescent signal was recorded. The difference in cell viability rates was found to be statistically significant. Data represents mean ± S.E.M. of eight independent experiments (n=8). ***p < 0.001, *p < 0.05, n.s.: non-significant.

Annexin V staining was used to assess whether CD47 siRNA-treated cells showed changes in apoptosis. No significant difference was observed between control and CD47 siRNA-treated cells (early apoptosis: 4.97 ± 0.727% versus 6.49 ± 1.59%, n.s., late-phase apoptosis: 0.283 ± 0.095% versus 0.320 ± 0.151%, n.s.) (Figures 14D, 14E).

We further examined the effect of CD47 downregulation on MDA-MB-468 viability: CD47 siRNA-treated cells showed significantly reduced viability compared to controls (74.8 ± 12.4% versus 99.8 ± 5.18%, p < 0.001) (Figure 14F).

#### 3.2.4 Reduced CD47 expression enhanced migration of MDA-MB-468 cells

We assessed the effect of CD47 knockdown on MDA-MB-468 migration using the wound healing assay. At 24 hours, the residual wound area was significantly smaller for CD47 siRNA-treated cells than controls (15.8 ± 2.70% versus 50.3 ± 16.7%, p < 0.05) (Figure 15A, 15B), though this difference became non-significant by 48 hours (4.06 ± 6.94% versus 13.6 ± 14.7%, n.s.). These results suggest CD47 knockdown produces a relatively acute increase in migratory capacity within the first 24 hours.

**Figure 15.**
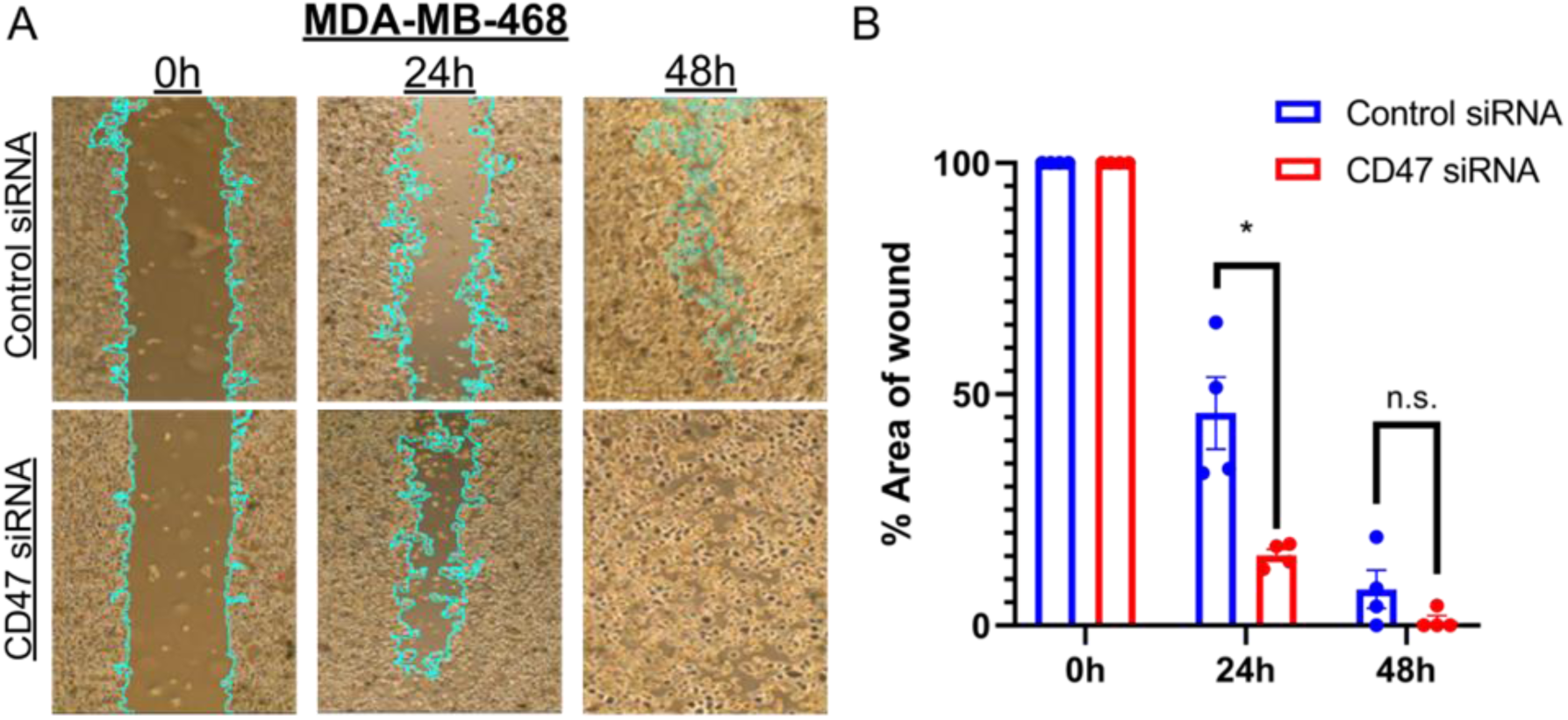
Effect of CD47 silencing on the migratory capability of MDA-MB-468 cells as demonstrated by wound healing assay. (A) Representative microscopy images of control siRNA-treated MDA-MB-468 cells (top row) and CD47 siRNA-treated MDA-MB-468 cells (bottom row) at 10x magnification at 0, 24 and 48 h after silicone inserts were removed, producing a 500 µm gap resembling a wound. (B) Bar graph depicting percentage area of wound for control siRNA-treated MDA-MB-468 cells and CD47 siRNA-treated MDA-MB-468 cells. Differences in the area of wound at 24 h between control siRNA-treated MDA-MB-468 cells and CD47 siRNA-treated MDA-MB-468 cells were significant. Results were normalised to the areas of the respective wounds at 0 h and represent the mean ± S.E.M. of four independent experiments (n=4). *p < 0.05, n.s.: non-significant

#### 3.2.5 Reduced CD47 expression in MDA-MB-468 cells conferred resistance against doxorubicin and oxaliplatin

We next assessed how CD47 expression affects MDA-MB-468 sensitivity to chemotherapeutic agents clinically used in TNBC. Dose-response analysis (Figure 16A–D) showed CD47 siRNA-treated cells had higher IC50 values than control siRNA-treated cells for doxorubicin (425 nM versus 183 nM) and oxaliplatin (22.4 μM versus 14.7 μM), and modestly lower IC50 values for paclitaxel (18.3 nM versus 23.6 nM) and docetaxel (8.77 nM versus 10.5 nM) (Figure 16).

**Figure 16.**
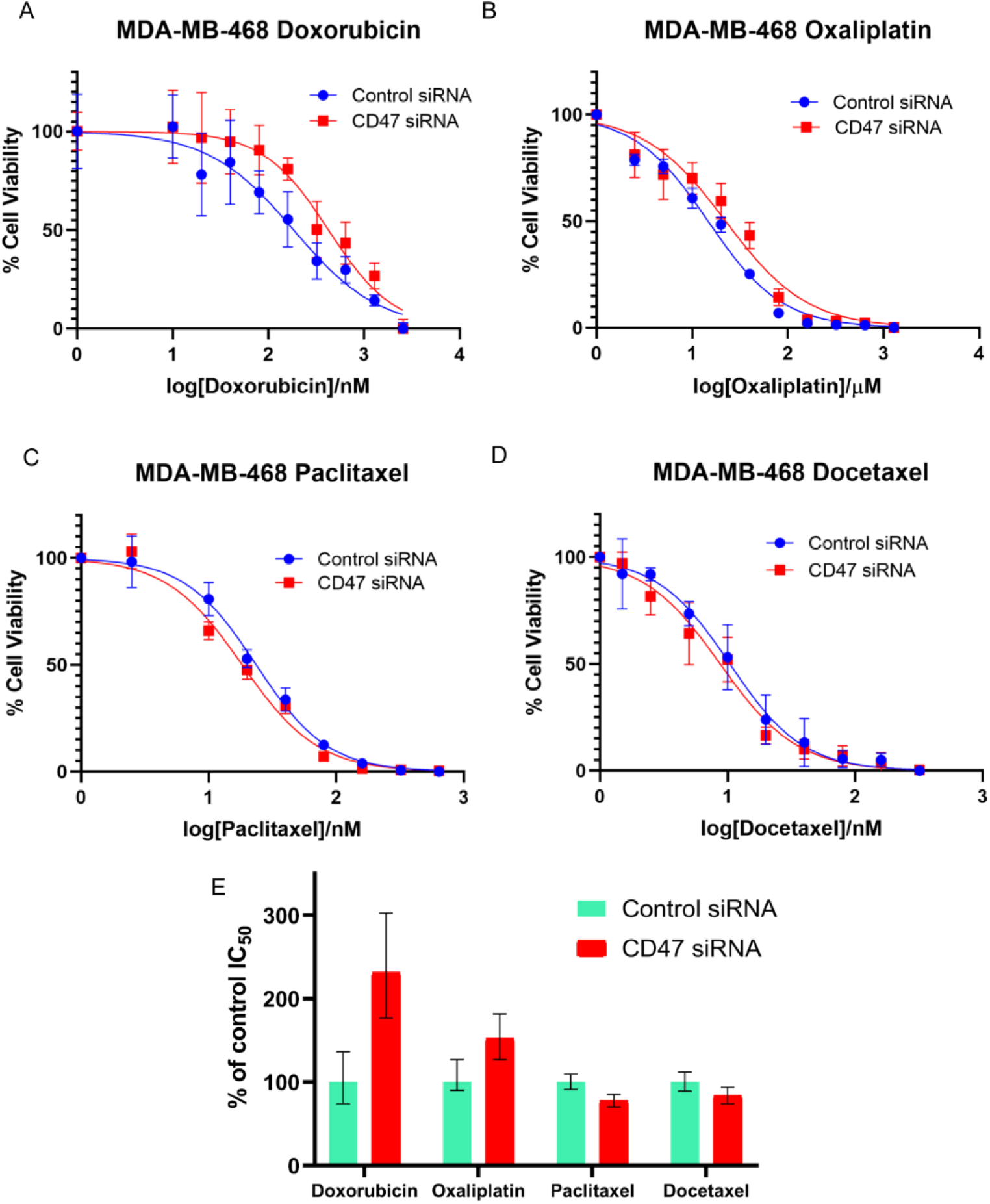
Dose-response relationships of viability of control siRNA versus CD47 siRNA-treated MDA-MB-468 cells upon treatment with doxorubicin, oxaliplatin, paclitaxel and docetaxel as determined via drug response assays. Dose response curves of control siRNA-treated and CD47 siRNA-treated MDA-MB-468 cells when treated with (A) doxorubicin, (B) oxaliplatin, (C) paclitaxel or (D) docetaxel. Error bars represent the S.D. of each data point. (E) Percentage difference in IC_50_ between control siRNA-treated and CD47 siRNA-treated MDA-MB-468 cells when treated with doxorubicin, oxaliplatin, paclitaxel or docetaxel. Results were normalised to the IC_50_ of control siRNA-treated MDA-MB-468 cells. Error bars represent the 95% confidence interval estimates of the IC_50_ as generated in GraphPad Prism 8.0.2.

## 4. Discussion

This study aimed to elucidate the non-canonical, non-immune biological effects of CD47 overexpression across different cancer types. A deeper understanding of these effects allows us to better assess the implications of CD47-targeted therapy for cancer patients and supports the rational design of synergistic treatment strategies that account for both the immune and non-immune functions of CD47. Immune evasion itself arises through diverse, often cancer type-specific mechanisms. For example, TP53 mutations disrupt T-cell, NK-cell, and macrophage-mediated tumour surveillance through distinct pathways^35^. This highlights the need to dissect the specific, non-immune contributions of individual checkpoints such as CD47 to overall tumour biology.

Our results suggest that CD47 exerts contrasting, context-dependent effects across cancer types: in our TNBC models, higher CD47 expression was associated with better cellular viability and proliferation, whereas in our DLBCL models, higher CD47 expression correlated with poorer survival outcomes. To explain this divergence, we explored the non-immunological roles CD47 plays in a solid tumour versus a haematological malignancy. Metabolic reprogramming is a hallmark of cancer, and metabolic alterations in DLBCL are closely linked to tumorigenesis, disease progression, and metastasis^21^. DLBCL is a metabolically active malignancy, characterised by increased activity of the mitochondrial electron transport chain and elevated expression of mitochondrial translation factors that together enhance oxidative phosphorylation^22^.

CD47 knockout reduced OCR in DLBCL cells, indicating mitochondrial dysfunction and a decline in respiration-driven ATP output. Because ECAR was unchanged in our DLBCL models, these cells appear primarily dependent on oxidative phosphorylation for cellular metabolism, with little reliance on aerobic glycolysis (the Warburg effect). The paradoxical increase in PGC1α expression following CD47 knockout may reflect a compensatory rise in mitochondrial biogenesis and activity^23^, potentially shifting DLBCL cells further toward oxidative phosphorylation rather than glycolysis. However, despite this apparent mitochondrial accumulation, both the JC-1 and Seahorse MitoStress assay results suggest that a substantial proportion of these mitochondria were unhealthy and dysfunctional. These metabolic and apoptotic vulnerabilities may also intersect with other non-phagocytic mechanisms of CD47 blockade: in lymphoid malignancies, CD47 blockade has been shown to trigger necroptotic cell death independent of its canonical anti-phagocytic function, complementing BCL-2-directed apoptosis induction^36^. This raises the possibility that CD47 loss sensitises DLBCL cells to R-CHOP through convergent, non-immune death pathways in addition to the mitochondrial dysfunction described here.

Although GAS8 remains poorly studied in cancer, it has been implicated in regulating the Hedgehog signalling cascade within the cilium^24^. Its presumed downregulation following CD47 knockout may therefore affect activation of this pathway, which is itself implicated in DLBCL pathogenesis: one study reported that Hedgehog pathway inhibition predominantly induced cell cycle arrest in GCB-subtype DLBCL cells and apoptosis in ABC-subtype cells^25^, with a corresponding reduction in cell viability. In our DLBCL models, CD47 knockout downregulated Shh, a key Hedgehog pathway regulator, suggesting CD47 involvement in this cascade. Shh has separately been shown to directly regulate mitochondrial function in neuronal cells^26^, so the relevance of CD47 influence on Shh and its downstream effects on mitochondria in DLBCL warrants further validation. ABCB1 (MDR1), an ATP-dependent efflux pump implicated in multidrug resistance across multiple cancer types^27^, is energetically costly to operate and is itself tied to mitochondrial function and metabolism: ABCB1 knockdown directly reduces mitochondrial ATP production^28^. In our study, the CD47-associated downregulation of ABCB1 could therefore have contributed to reduced mitochondrial activity in DLBCL, and may also lower the likelihood of R-CHOP multidrug resistance, which is consistent with a role for CD47 inhibition in improving drug sensitivity. These directional hypotheses will need to be confirmed experimentally, for instance through ABCB1 overexpression.

Notably, although our initial CD47 knockout DLBCL clones lacked detectable cell-surface CD47, western blotting revealed that CD47 persisted intracellularly in some subsequent clones. These clones subsequently regained partial cell-surface CD47 expression on flow cytometry, indicating that they represented a knockdown rather than a true knockout phenotype.

In TNBC, CD47 downregulation significantly delayed cell cycle progression in the G1 phase. Because cell cycle arrest is closely linked to both apoptosis and proliferation, and a greater proportion of CD47-silenced cells appeared enlarged and flattened, which was consistent with G1-phase cells expanding as proteins and organelles are synthesised in preparation for mitosis^29^. CD47 knockdown significantly reduced expression of CCND1, a plausible driver of the observed cell cycle arrest. Flow cytometric analysis of phosphatidylserine externalisation and membrane integrity showed no difference in apoptotic proportions after CD47 knockdown, indicating that G1-arrested, CD47-silenced cells did not undergo mitotic catastrophe. We therefore asked whether the rate of cell division itself had slowed: cell viability assays showed significantly fewer CD47-knockdown cells than controls after 48 hours, consistent with reduced proliferation, either because cells took longer to traverse G1 or because a subset became senescent, where cells remained metabolically active but no longer proliferative. Together, these findings suggest that TNBC cells with relatively higher CD47 expression retain greater viability and proliferative capacity, consistent with clinical studies linking increased CD47 expression to increased tumour growth and poorer prognosis in TNBC patients^16,30^. Paradoxically, this contrasts with studies reporting that CD47 knockdown reduces migration and invasion in other cancer types^31,32^, as our data instead showed enhanced migration. A plausible explanation is that upregulation of migration- and invasion-related genes represents a compensatory feedback response to the abrupt loss of CD47 in an intrinsically high-CD47-expressing cell line, potentially reflecting a role for CD47 in maintaining cellular homeostasis^33^. CD47 silencing also selectively altered chemosensitivity in MDA-MB-468 cells. Testing doxorubicin, paclitaxel, docetaxel, and oxaliplatin, which represents the anthracycline, taxane, and platinum-based regimens used clinically in TNBC, we found that CD47 knockdown conferred resistance to doxorubicin and oxaliplatin while modestly enhancing sensitivity to the taxanes. This pattern is consistent with a report that FASN overexpression in breast cancer cells drives resistance to DNA-damaging agents such as doxorubicin and cisplatin while sparing microtubule-stabilising agents like paclitaxel^34^. As CD47 silencing has separately been linked to enhanced FASN expression, this may explain the selective resistance to doxorubicin and oxaliplatin, but not the taxanes, that we observed after CD47 knockdown in MDA-MB-468 cells. More broadly, breast cancer cells have been shown to adopt cooperative, altruistic behaviours, including paracrine signalling that confers taxane resistance on neighbouring cells at a fitness cost to the signalling cell itself. This suggests that the chemoresistance phenotypes we observed after CD47 knockdown may also be shaped by similar cell-cell cooperative dynamics within the tumour population^37,38^.

This study has several limitations. First, our findings rely on *in vitro* cell line models with CD47 overexpression well above typical physiological levels. *In vivo* validation using mouse models will be necessary to confirm that these effects hold at physiologically relevant CD47 levels and translate clinically. Second, we plan to complement our knockdown/knockout approach with the inverse experiment: overexpressing CD47 in relatively low-CD47-expressing lines such as MDA-MB-453 or SUDHL2, to extrapolate our findings across the full spectrum of CD47 expression levels observed in cancer. For stable, high-efficiency CD47 overexpression, we anticipate using lentiviral transduction protocols optimised for primary and immortalised cell models^39^, which would complement the transient plasmid-based approach used in our current work.

Importantly, CD47-associated R-CHOP sensitivity in our DLBCL models was dependent on the presence of human serum, pointing to a possible role for CD47 in complement-dependent cytotoxicity (CDC) that warrants further investigation. Collectively, our data suggest CD47 influences R-CHOP sensitivity in DLBCL through both intrinsic (mitochondrial metabolic health) and extrinsic (CDC) mechanisms, presenting opportunities to augment first-line R-CHOP efficacy. In TNBC, our results suggest CD47 expression could help stratify patients toward optimal treatment regimens: anthracycline- or platinum-based regimens may be more effective in higher-CD47-expressing tumours, while taxane-based regimens, or combination with CD47-downregulating agents, may better suit tumours with intrinsically low CD47 expression.

## 5. Conclusion

In conclusion, this study provides insight into the non-canonical, non-immune biological effects of varying CD47 expression across cancer types. In TNBC, CD47 silencing delayed cell cycle progression, likely underlying the reduced viability we observed, while also driving chemoresistance to DNA-damaging agents and enhanced sensitivity to taxanes. In DLBCL, CD47 silencing induced mitochondrial dysfunction, impaired oxidative phosphorylation, and sensitised cells to first-line R-CHOP therapy in the presence of human serum. Together, these findings offer a rational basis for stratifying chemotherapeutic drug selection by tumour CD47 expression, while also serving as a caveat for the clinical use of CD47-targeted agents: because CD47 exerts contrasting, cancer type-specific effects beyond immune evasion, further validation studies are needed to define its full biological repertoire before these agents are deployed in combination regimens. A more complete understanding of CD47-mediated signalling in cancer cells will be essential to designing effective combination strategies that pair CD47 downregulation or blockade with existing therapies to improve clinical outcomes.

## Ethics Approval and Consent to Participate

FFPE patient samples from patients were obtained from the Department of Pathology, National University of Singapore, with prior approval obtained from NUS IRB.

## Author Contribution Statement

S.S.S.H. and M.S.B.M. conceptualized the study, provided funding and administrative oversight. S.M.L., M.S.B.M., T.C.I.L. and J.Y.M.T. provided intellectual discussion, designed and performed experiments. T.C.I.L. and J.Y.M.T. analysed the data and wrote the manuscript. F.J.H.N. designed and conducted experiments.

## Acknowledgements

We thank Ms Wang Xiaoning of the Flow Cytometry Laboratory Unit of the Yong Loo Lin School of Medicine, National University of Singapore, for flow cytometry support. S.M.L. and M.S.B.M. acknowledge the NUHS Seed Grant (NR21MRF226OM) from the Department of Pathology, National University of Singapore. S.M.L, M.S.B.M. and J.Y.M.T. acknowledge funding from the N2CR SCOUT Pilot Grant Call 2026. M.S.B.M. acknowledges support from the NUS Postgraduate Research Scholarship and NUSMed Postdoctoral Fellowship. J.Y.M.T acknowledges support from the NUS Singapore Graduate Fellowship/Graduate Tutor scheme.

## List of Abbreviations

ABC: Activated B-cell
CD47: Cluster of Differentiation 47
DLBCL: Diffuse large B-cell lymphoma
ECAR: Extracellular acidification rate
GCB: Germinal center B-cell
NHL: non-Hodgkin lymphoma
NUH: National University Hospital, Singapore
OCR: Oxygen consumption rate
siRNA: Small interfering ribonucleic acid
TNBC: Triple negative breast cancer

## References

1. Tan, D., Chan, J. Y., Wudhikarn, K., Wong, R. S. M., Poon, L., Norasetthada, L., Huang, T. C., & Tse, E. (2024). Unmet Needs in the First-Line Treatment of Diffuse Large B-cell Lymphoma: Expert Recommendations From the Asia-Pacific Region With a Focus on the Challenging Subtypes. Clinical lymphoma, myeloma & leukemia, 24(9), e320–e328. 10.1016/j.clml.2024.05.013

2. Schneider C, Pasqualucci L, Dalla-Favera R. (2013). Molecular pathogenesis of diffuse large B-cell lymphoma. Semin Diagn Pathol. 28(2):167–77. doi: 10.1053/j.semdp.2011.04.001.

3. Mesin L, Ersching J, Victora GD. (2016). Germinal Center B Cell Dynamics. Immunity. 45(3):471–482. doi: 10.1016/j.immuni.2016.09.001.

4. Singapore Cancer Registry Annual Report. (2022). https://nrdo.gov.sg/publications/cancer

5. Gumà J, Palazón-Carrión N, Rueda-Domínguez A, Sequero S, Calvo V, García-Arroyo R, Gómez-Codina J, Llanos M, Martínez-Banaclocha N, Provencio M. (2025). SEOM-GOTEL clinical guidelines on diffuse large B-cell lymphoma (update 2025). Clin Transl Oncol. 27(12):4381–4392. doi: 10.1007/s12094-025-04080-z.

6. Liu Y, Barta SK. (2019). Diffuse large B-cell lymphoma: 2019 update on diagnosis, risk stratification, and treatment. Am J Hematol. 94(5):604–616. doi: 10.1002/ajh.25460.

7. Yin, L., Duan, J. J., Bian, X. W., and Yu, S. C. (2020). Triple-negative breast cancer molecular subtyping and treatment progress. Breast cancer research : BCR, 22(1), 61

8. Xiong N, Wu H and Yu Z. (2024). Advancements and challenges in triple-negative breast cancer: a comprehensive review of therapeutic and diagnostic strategies. Front. Oncol. 14:1405491. doi: 10.3389/fonc.2024.140549

9. Wu, Q., Siddharth, S., and Sharma, D. (2021). triple-negative Breast Cancer: A Mountain Yet to Be Scaled Despite the Triumphs. Cancers, 13(15), 3697.

10. Si, Y., Zhang, Y., Guan, J. S., Ngo, H. G., Totoro, A., Singh, A. P., Chen, K., Xu, Y., Yang, E. S., Zhou, L., Liu, R., and Liu, X. M. (2021). Anti-CD47 Monoclonal Antibody-Drug Conjugate: A Targeted Therapy to Treat Triple-Negative Breast Cancers. Vaccines, 9(8), 882.

11. Yao, H., He, G., Yan, S., Chen, C., Song, L., Rosol, T. J., and Deng, X. (2017). Triple-negative breast cancer: is there a treatment on the horizon?. Oncotarget, 8(1), 1913–1924

12. Uchimiak K, Badowska-Kozakiewicz AM, Sobiborowicz-Sadowska A, Deptała A. (2022). Current State of Knowledge on the Immune Checkpoint Inhibitors in Triple-Negative Breast Cancer Treatment: Approaches, Efficacy, and Challenges. Clin Med Insights Oncol. 16:11795549221099869. doi: 10.1177/11795549221099869

13. Chao MP, Weissman IL, Majeti R. (2012). The CD47–SIRPα pathway in cancer immune evasion and potential therapeutic implications. Current Opinion in Immunology. 24(2):225–32

14. Chen C, Wang R, Chen X, Hou Y, Jiang J. (2022). Targeting CD47 as a Novel Immunotherapy for Breast Cancer. Front Oncol. 12:924740. doi: 10.3389/fonc.2022.924740.

15. Chao MP, Alizadeh AA, Tang C, Myklebust JH, Varghese B, Gill S, et al. (2010). Anti-CD47 Antibody Synergizes with Rituximab to Promote Phagocytosis and Eradicate Non-Hodgkin Lymphoma. Cell. 142(5):699–713

16. Yuan J, Shi X, Chen C, He H, Liu L, Wu J, Yan H. (2019). High expression of CD47 in triple negative breast cancer is associated with epithelial-mesenchymal transition and poor prognosis. Oncol Lett. 18(3):3249–3255. doi: 10.3892/ol.2019.10618.

17. Frazier EP, Isenberg JS, Shiva S, Zhao L, Schlesinger P, Dimitry J, et al. (2011). Age-dependent regulation of skeletal muscle mitochondria by the thrombospondin-1 receptor CD47. Matrix Biology. 30(2):154–61

18. Hu T, Liu H, Liang Z, Wang F, Zhou C, Zheng X, et al. (2020). Tumor-intrinsic CD47 signal regulates glycolysis and promotes colorectal cancer cell growth and metastasis. Theranostics. 10(9):4056–72

19. Chiche J, Reverso-Meinietti J, Mouchotte A, Rubio-Patiño C, Mhaidly R, Villa E, et al. (2019). GAPDH Expression Predicts the Response to R-CHOP, the Tumor Metabolic Status, and the Response of DLBCL Patients to Metabolic Inhibitors. Cell Metabolism. 29(6):1243–57.e10

20. Zhang, M.Q., Lan, J.S., Li, Z., Yang, S.Q., Gu, D.H., Nie, W.L., Ding, Y., and Zhang, T. (2025). Ferroptosis meets biomimetic nano-systems: A novel strategy for targeted cancer therapy. Cell Biomater. 1, 100023. 10.1016/j.celbio.2025.100023.

21. Pang, Y., Lu, T., Xu-Monette, Z. Y., & Young, K. H. (2023). Metabolic Reprogramming and Potential Therapeutic Targets in Lymphoma. International Journal of Molecular Sciences, 24(6), 5493. 10.3390/ijms24065493

22. Norberg E, Lako A, Chen P.H, Stanley I.A, Zhou F, Ficarro S.B, Chapuy B, Chen L, Rodig S, Shin D, et al. (2017). Differential contribution of the mitochondrial translation pathway to the survival of diffuse large B-cell lymphoma subsets. Cell Death Differ, 24: 251–262.

23. Wang, Y., Peng, J., Yang, D., Xing, Z., Jiang, B., Ding, X., Jiang, C., Ouyang, B., & Su, L. (2024). From metabolism to malignancy: the multifaceted role of PGC1α in cancer. Frontiers in oncology, 14, 1383809. 10.3389/fonc.2024.1383809

24. Lewis WR, Malarkey EB, Tritschler D, Bower R, Pasek RC, et al. (2016). Mutation of Growth Arrest Specific 8 Reveals a Role in Motile Cilia Function and Human Disease. PLOS Genetics, 12(7): e1006220. 10.1371/journal.pgen.1006220

25. Singh, R., Kim, J., Davuluri, Y. et al. (2010). Hedgehog signaling pathway is activated in diffuse large B-cell lymphoma and contributes to tumor cell survival and proliferation. Leukemia, 24, 1025–1036. 10.1038/leu.2010.35

26. Malhotra, A., Dey, A., Prasad, N., & Kenney, A. M. (2016). Sonic Hedgehog Signaling Drives Mitochondrial Fragmentation by Suppressing Mitofusins in Cerebellar Granule Neuron Precursors and Medulloblastoma. Molecular cancer research : MCR, 14(1), 114–124. 10.1158/1541-7786.MCR-15-0278.

27. Skinner, K. T., Palkar, A. M., & Hong, A. L. (2023). Genetics of *ABCB1* in Cancer. Cancers, 15(17), 4236. 10.3390/cancers15174236.

28. Giddings, E.L., Champagne, D.P., Wu, MH. et al. (2021). Mitochondrial ATP fuels ABC transporter-mediated drug efflux in cancer chemoresistance. Nat Commun, 12, 2804. 10.1038/s41467-021-23071-6

29. Sommer, R. A., DeWitt, J. T., Tan, R., and Kellogg, D. R. (2021). Growth-dependent signals drive an increase in early G1 cyclin concentration to link cell cycle entry with cell growth. eLife, 10, e64364.

30. Zhang, H., Lu, H., Xiang, L., Bullen, J. W., Zhang, C., Samanta, D., Gilkes, D. M., He, J., and Semenza, G. L. (2015). HIF-1 regulates CD47 expression in breast cancer cells to promote evasion of phagocytosis and maintenance of cancer stem cells. Proceedings of the National Academy of Sciences of the United States of America, 112(45), E6215–E6223.

31. Zhao, H., Wang, J., Kong, X., Li, E., Liu, Y., Du, X., Kang, Z., Tang, Y., Kuang, Y., Yang, Z., Zhou, Y., and Wang, Q. (2016). CD47 Promotes Tumor Invasion and Metastasis in Non-small Cell Lung Cancer. Scientific reports, 6, 29719.

32. Liu, Y., Chang, Y., He, X., Cai, Y., Jiang, H., Jia, R., and Leng, J. (2020). CD47 Enhances Cell Viability and Migration Ability but Inhibits Apoptosis in Endometrial Carcinoma Cells via the PI3K/Akt/mTOR Signaling Pathway. Frontiers in oncology, 10, 1525.

33. El-Brolosy, M. A., & Stainier, D. Y. R. (2017). Genetic compensation: A phenomenon in search of mechanisms. PLoS genetics, 13(7), e1006780. 10.1371/journal.pgen.1006780

34. Liu, H., Wu, X., Dong, Z., Luo, Z., Zhao, Z., Xu, Y., and Zhang, J. T. (2013). Fatty acid synthase causes drug resistance by inhibiting TNF-α and ceramide production. Journal of lipid research, 54(3), 776–785.

35. Wang, C., Tan, J. Y. M., Chitkara, N., and Bhatt, S. (2024). TP53 Mutation-Mediated Immune Evasion in Cancer: Mechanisms and Therapeutic Implications. Cancers, 16(17), 3069. 10.3390/cancers16173069

36. Chong, S. J. F., Valentin, R., Wang, J., Zhu, F., Gokhale, P. C., Eschle, B. K., et al. (2026). CD47 blockade-driven necroptosis complements BCL-2 inhibition-driven apoptosis in lymphoid malignancies. Journal of Hematology & Oncology, 19(1), 11. 10.1186/s13045-025-01774-3

37. Masroni, M. S. B., Lee, K. W., Lee, V. K. M., Ng, S. B., Law, C. T., Poon, K. S., et al. (2023). Dynamic altruistic cooperation within breast tumors. Molecular Cancer, 22, 206. 10.1186/s12943-023-01896-7

38. Masroni, M. S. B., Koay, E. S., Lee, V. K. M., Ng, S. B., Tan, S. Y., Tan, K. M., Archetti, M., and Leong, S. M. (2025). Sociobiology meets oncology: unraveling altruistic cooperation in cancer cells and its implications. Experimental & Molecular Medicine, 57(1), 30–40. 10.1038/s12276-024-01387-9

39. Tan, J. Y. M., Tan, J. C., Wang, C., Wu, L., Gascoigne, N. R. J., and Bhatt, S. (2025). Protocol for the simultaneous activation and lentiviral transduction of primary human T cells with artificial T cell receptors. STAR Protocols, 6(1), 103685. 10.1016/j.xpro.2025.103685

